# Developmentally-specific physiological and metabolic responses support drought resilience in switchgrass and constrains biofuel yield

**DOI:** 10.1101/2025.05.29.656847

**Authors:** Binod Basyal, Xingxing Li, V. J. Pargulski, Xinyu Fu, Nicole M. Nightingale, Katherine A. Overmyer, Joshua J. Coon, Yaoping Zhang, Gian Maria Niccolò Benucci, Robert L. Last, Trey K. Sato, Berkley J. Walker

**Author notes:** Deceased (Yaoping Zhang).

## Abstract

Switchgrass (*Panicum virgatum*) is a promising bioenergy crop due in part to its resilience to drought stress. However, the significance of drought timing remains poorly understood, both from a plant biology perspective and its impact on downstream biofuel production. This study determines the developmental stage-specific physiological and metabolic responses of switchgrass to drought stress and its implications for biofuel production using a custom-built programmable irrigation system. Vegetative, flowering, and senescence-stage drought significantly reduced CO_2_ assimilation, and stomatal conductance without affecting biomass yield. Metabolic profiling revealed significant accumulation of glucose, fructose, quinic acid, shikimate and GABA during vegetative-stage drought, while flowering and senescence stages exhibited limited metabolic changes. Similarly, specialized metabolites also displayed distinct developmental patterns, with vegetative-stage drought driving the most pronounced metabolic alterations. Thermochemically-treated and hydrolyzed switchgrass biomass from vegetative-stage drought showed elevated lignocellulose-derived compounds and saponins with the latter most positively correlating with fermentation lag times. Conversely, senescence-stage drought enhanced ethanol yields while lowering saponin levels in the hydrolysates. While vegetative-stage drought enhanced physiological resilience, it compromises downstream biofuel production by introducing fermentation inhibitors, particularly saponins.

## Introduction

Switchgrass (*Panicum virgatum* L.), a perennial C4 grass native to North America, is a promising bioenergy crop due to its resilience and yield even in marginal soils. As drought frequency and severity are predicted to increase with climate change^1^, it is critical to understand how crops like switchgrass respond to drought. While drought tolerance mechanisms in annual plants are well-studied, less is known about perennials which allocate resources differently across complex life cycles ^2^.

Drought stress reduces photosynthesis, biomass production and alters metabolism ^3,4^. The effects of drought can vary across developmental stages due to changes in carbon allocation, such as the storage of carbohydrates in rhizomes. While there is work investigating the metabolic response to a single drought imposed during the vegetative stage, there is limited information on how switchgrass metabolite profiles shift with drought across these stages. A major knowledge gap lies in the distinction between core physiological and metabolic responses to drought and developmentally specific physiological and metabolic changes. While switchgrass exhibits a range of metabolic adjustments in response to environmental stresses, it remains unclear which of these responses are fundamental, conserved pathways activated across all developmental stages, and which are unique to specific growth phases such as tillering, reproduction, or senescence.

Another critical gap is understanding how drought timing affects biomass composition, which consequently affects biofuel yield and quality. Drought can alter biomass composition, resulting in production of inhibitory compounds in the raw biomass and during the pretreatment process that hamper fermentation^5^. For example, biomass from drought years can inhibit fermentation^6^, initially attributed to higher soluble sugars reacting with ammonia during pretreatment, forming fermentation inhibitors like imidazoles and pyrazines. Additionally, specialized metabolites such as saponins produced under drought^8^ as antimicrobial or anti-herbivory compounds have also been shown to strongly inhibit fermentation^8^. Alternatively, drought can increase levels of lignocellulose derived inhibitors like ferulic acid and *p*-coumaric acid^5^, which can further degrade into toxic amides during the pretreatment process which could inhibit microbial fermentation^9^. While these are several putative inhibitors to biofuel production, the timing of the production of these inhibitors in response to drought is unclear. Furthermore, how the production and accumulation of non-lignocellulosic specialized metabolites potentially toxic to *Saccharomyces cerevisiae*, such as saponins, change in response to drought stress has not been thoroughly studied. To the best of our knowledge, this study provides the first direct identification of saponins in the hydrolysate samples.

In this study, switchgrass was grown from seedling stage through senescence and subjected to four watering treatments: a continuously well-watered control and drought imposed at specific developmental stages (vegetative, flowering, and senescence). This single-factor experimental design with developmental stage-specific drought aims to address the following critical questions (1) How does switchgrass metabolically respond to drought stress? (2) Are these responses dependent on the developmental stage at which drought occurs? (3) Do these metabolic shifts influence biomass fermentation? (4) Among the array of drought-induced compounds, which metabolites most strongly inhibit fermentation? By examining these questions, the study seeks to fill key knowledge gaps in switchgrass drought responses and how these responses impact biofuel yield and quality.

## Materials and Methods

### Plant Material and Growth Conditions

Switchgrass (*Panicum virgatum L.*), cv. ‘Cave-In-Rock,’ was cultivated in a greenhouse at Michigan State University under controlled temperature (Averaged 26.6°C) and light conditions (16-hour photoperiod). The plants were initially grown from seeds that were germinated on wet filter paper placed over vermiculite medium in petri dishes. A week after emergence, the seedlings were transplanted into 8 L pots containing potting mix (Michigan Grower Products, Inc, Michigan, USA, Supplementary table 1). Prior to the drought treatment, all plants were watered to an ambient soil moisture of 25% Volumetric Water Content (VWC).

### Drought Treatment and Experimental Design

Plants were assigned to four treatment groups, vegetative drought, flowering drought, senescence drought and a well-watered control. Following a 30-day establishment period under ambient watering, drought was imposed by withholding irrigation until soil VWC reached 1% (supplementary fig. 1), which was then maintained volumetrically to elicit physiological drought responses. Control plants and plants not included in the particular drought treatment were maintained at 25% VWC using an automated irrigation system controlled using Raspberry Pi computers and TEROS 12 moisture probes (METER Group, Inc. USA). Drought treatments were applied only during the designated developmental stages 33 days (vegetative), 39 days (flowering), and 20 days (senescence). All treatments were randomly assigned within three blocks, and the experiment spanned 122 days.

### Gas Exchange and Chlorophyll Fluorescence Measurements

Gas exchange and chlorophyll fluorescence parameters, including stomatal conductance (gₛ), net CO₂ assimilation (A), and the operating efficiency of photosystem II (ΦPSII), were measured as survey measurement ^35^ using a LI-COR 6800 portable photosynthesis system (LI-COR Biosciences, Lincoln, NE). The chamber environment was controlled to greenhouse conditions for CO_2_ concentration, light, temperature and vapor pressure deficit, for each survey day. The reference CO₂ concentration was set at 400 ppm, and the chamber area of 6 cm² was corrected for blade width. Photosynthetically Active Radiation was measured using a PAR sensor before each measurement. All gas exchange measurements for that day were done at that constant PAR. Similarly, a constant temperature exchange and vapor pressure deficit value were used for all the measurements for each survey day. Measurements were taken at 14 different time points throughout the experimental duration.

### Biomass and Growth Measurements

Tillering and the number of green leaves were recorded throughout the experiment to assess growth responses to drought stress. At the end of the experiment, both aboveground and belowground biomass was harvested, dried at 65°C for 72 hours, and weighed.

### Cell Wall Composition analysis

Total lignin content was quantified using the acetyl bromide soluble lignin (ABSL) assay. Approximately 2 mg (±0.5 mg) of alcohol-insoluble residue (AIR) was weighed into 2 mL microcentrifuge tubes, including three biomass controls and two blanks. Following acetone rinsing and overnight drying, 250 μL of 25% acetyl bromide in glacial acetic acid was added. Samples were incubated at 50°C for 3 h, cooled, centrifuged at 10,000 RPM, and 100 μL of supernatant was mixed with hydroxylamine hydrochloride, NaOH, and acetic acid to a final volume of 2.0 mL. Absorbance was read at 280 nm.

Lignin composition was assessed using thioacidolysis followed by TMS derivatization. AIR (2 mg) was reacted with a BF₃/EtSH/dioxane mixture, sealed under nitrogen, and heated at 100°C for 4 h. The product was neutralized, extracted with ethyl acetate, derivatized with pyridine and N,O-bis(trimethylsilyl)acetamide, and analyzed by GC.

Matrix polysaccharides were analyzed by sequential washing of dried biomass, starch digestion using amylase and pullulanase, and trifluoroacetic acid hydrolysis. Alditol acetates were extracted for GC analysis. Crystalline cellulose was measured using the Updegraff method and sulfuric acid hydrolysis.

Starch content was quantified via enzymatic digestion with α-amylase and amyloglucosidase followed by glucose measurement using a glucose oxidase/peroxidase calorimetric assay.

### Metabolite Profiling

#### Central metabolites

Frozen tissues were homogenized cryogenically using a FastPrep-24 instrument with a CoolPrep Adapter and dry ice. Metabolites were extracted with methanol:chloroform:water, spiked with ribitol as an internal standard. After centrifugation, the polar phase was lyophilized and derivatized with methoxyamine in pyridine followed by trimethylsilylation. A 1 µL aliquot was analyzed using an Agilent 7890A GC coupled to a 5975C MS. Oven temperature was ramped from 40°C to 320°C. Mass spectra (m/z 50–600) were acquired via electron impact ionization. Metabolites were identified against the NIST 14 Library and standards; relative quantification was based on normalized peak areas.

#### Metabolite extraction for untargeted LC-MS analysis

Total metabolite extraction from fresh switchgrass biomass followed a published protocol^36^. Tissues were flash-frozen in liquid nitrogen, ground, and incubated overnight at 4°C in 80% methanol with 0.5 µM telmisartan and 0.1% formic acid. Supernatants were collected post-centrifugation and stored at –20°C for LC-MS. For hydrolysates, 500 µL was passed through C18 SPE cartridges. Flowthrough and methanol eluates were collected, dried, redissolved in extraction solvent, and transferred to HPLC vials. All samples were stored at –20°C prior to analysis.

#### LC-MS metabolomics analysis

The metabolite separation and MS were conducted using UPLC BEH C18 column (Waters, Milford, MA) and a Thermo Q-Exactive mass spectrometer (Thermo). The UPLC flow rate was 0.4 ml/min at 40 C. Each sample injection was 10 µl. The mass spectrometer operated at 35,000 a.u. resolution, sheath gas flow rate 50 a.u., aux gas flow rate 13 a.u., spray voltage 3.5 kV, capillary temp. 50 C and aux gas heater temp. 425 C. Positive ionization data dependent acquisition (DDA) scanned m/z 100 to 1500, selecting top five ions per scan for fragmentation

The raw mass spectral data in Thermo RAW format were transformed to mzML format using proteowizard (version 3). The mzML data were imported to MZmine3^37^ to identify the metabolite features based on retention time and mass-to-charge ratio (m/z) values. The resultant feature abundance table analyzed in MetaboAnalyst for relative quantification and statistical analysis. Abundances were normalized to an internal standard, log-transformed and “pareto”.The log-transformed and pareto scaled (normalized) abundances of features were then used for statistics. The feature annotation was performed by searching MassBank using Global Natural Products Social Molecular Networking and predicting chemical classes with CANOPUS (class assignment and ontology prediction using mass spectrometry in SIRIUS 4.

### Fermentation Assay and hydrolysate metabolomics

#### Biomass pretreatment and enzymatic hydrolysis

Dried and ground switchgrass samples were pretreated using a modified Soaking in Aqueous Ammonia (SAA) method^38^. 15 g of 2mm milled switchgrass was transferred into a glass pressure tube with 60 ml of 18% v/v diluted ammonium hydroxide. The tubes were sealed, inverted, and incubated at 75 °C, mixed manually after 1.5 hours, and incubated for another 2 hours. After incubation, the samples were dried under compressed air for 48 hand air driedfor 5 more days.

For hydrolysate production, dried SAA pretreated switchgrass was loaded into centrifuge tubes at 7% glucan loading in 50 ml total volume. After autoclaving with double distilled water and potassium phosphate, HCL was added, followed by sterile-filtered cellulase and xylanase enzymes. Tubes were incubated at 50C for 2 days, reloaded with enzymes and water, and incubated for 5 additional days. Hydrolysates were then centrifuged, pH adjusted to 5.8 and sterile-filtered.

#### Fermentation

Fermentations were carried out as previously described^39^ with modifications. Hydrolysates were diluted 3 parts hydrolysate:1 part sterile ddH_2_O. The yeast strain GLBRCY1455, engineered by deleting *FLO8* from GLBRCY1327^40^ was used. Measurements of cell growth, CO_2_ and ethanol production are described previously^39^. Yeast cultures were grown anaerobically, centrifuged during log-phase growth, resuspended and inoculated into 60 ml Wheaton serum bottles. Fermentation occurred at 30 °C. Gas production was measured using an AER-800 respirometer, and post-fermentation supernatants were analyzed by HPLC-refractive index detection (RID) for sugar and ethanol concentrations^41^. Final cell density (OD_600_) was measured with a Beckman DU720 spectrophotometer.

#### Hydrolysate Metabolomics

Samples were stored at 4°C, diluted 10x and 75x using using 2.5 µM 13C6 vanillin in waterand centrifuged to remove particulates. Supernatant was analyzed by LC-MS/MS. Tareted standards from 0.024 μM to 50 μM were used for quantification. Discovery metabolomics used 10x diluted samples.

For targeted lignotoxins assay, 5 μl sample was injected by Vanquish LC system and separated using an ACQUITY UPLC HSS T3 column at 40°C. Mobile phase A consisted of 0.1% formic acid in water. Mobile phase B consisted of 100% acetonitrile. A gradient of 0.1% formic acid (A) and 100% acetonitrile (B) was used. The LC was coupled to a Q Exactive Orbitrap mass spectrometer operating at polarity switching mode, acquiring full MS and targeted MS2 spectra. Source conditions included 275°C capillary temperature, 30 sheath gas units, and ±4.0–4.5 kV spray voltage. Data were manually processed in TraceFinder 4.0 using a 10 ppm mass tolerance, and summary statistics were exported to Excel.

### Statistical Analysis

Data were analyzed using R software (R Core Team, 2024). Two-way repeated measures ANOVA evaluated the effects of drought timing, survey time, and their interaction on tillering, leaf number, stomatal conductance, net CO₂ assimilation, and Φ_PSII_. One-way ANOVA assessed biomass yield, lignin content, ethanol production, glucose consumption, and yeast OD₆₀₀ followed by Tukey’s (p<0.05).

Fermentation lag time and rates were determined by fitting the CO₂ evolution data to the sigmoidal model:

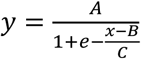

where A = maximum value of the curve (upper bound) x = time

e = Euler’s number

B = inflection point, where the curve changes shape

C = rate/steepness of the curve.

One-way ANOVA and Tukey’s test evaluated metabolite levels, with a false discovery rate threshold of 5%. Principal component analysis (PCA) and heatmaps were used to explore metabolic differences between treatments and developmental stages. A significance level of p < 0.05 was applied for all tests.

Linear mixed-effect models predicted fermentation lag time from saponin content which was scaled and centered on the mean [i.e. x-µ/sd(x)]. Models with fixed and random slopes were compared. R^2^ Values, diagnostics, and estimates are available (supplementary fig. 12-14, supplementary table 3).

## Results

### Drought impairs switchgrass physiology across developmental stages

Drought stress was applied at each developmental stage, while all other stages remained well-watered. The control group was well-watered throughout all stages. For drought treatments, soil moisture was reduced to 1% volumetric water content (VWC) during one each of the developmental stages investigated (vegetative, flowering, and senescence, Supplementary fig. 1). When a physiological drought response was determined, the plants were re-watered to 25% VWC until the next developmental stage was reached by the control and non-droughted treatments. The effectiveness of the drought was confirmed by a significant reduction in leaf water potential (LWP, Supplementary fig. 2). The most severe reduction in LWP occurred during senescence stage where it was 2.5 times lower in droughted plants compared to the control. Gas exchange and chlorophyll fluorescence were measured to assess photosynthetic and physiological changes to drought timing. Net CO₂ assimilation significantly declined during peak drought at each stage, with the most substantial drop occurring during the vegetative stage, 53 days after transplantation (fig. 1 A). Drought during the vegetative stage decreased net CO₂ assimilation by 35% and stomatal conductance by 49% (fig. 1 B). Chlorophyll fluorescence measurement indicated a decrease in operating efficiency of PSII (ɸ_PS2_) by approximately 17%, 25%, and 36% during the vegetative, flowering, and senescence stages, respectively (fig. 1 C). Seasonal declines in net CO₂ assimilation and ɸ_PSII_ typical of perennial grasses were observed across all treatments, with the most pronounced decrease in the senescence drought treatment (68% vs. 89% for CO₂ assimilation and 57% vs. 74% for ɸ_PSII_, fig. 1 A).

**Fig. 1.**
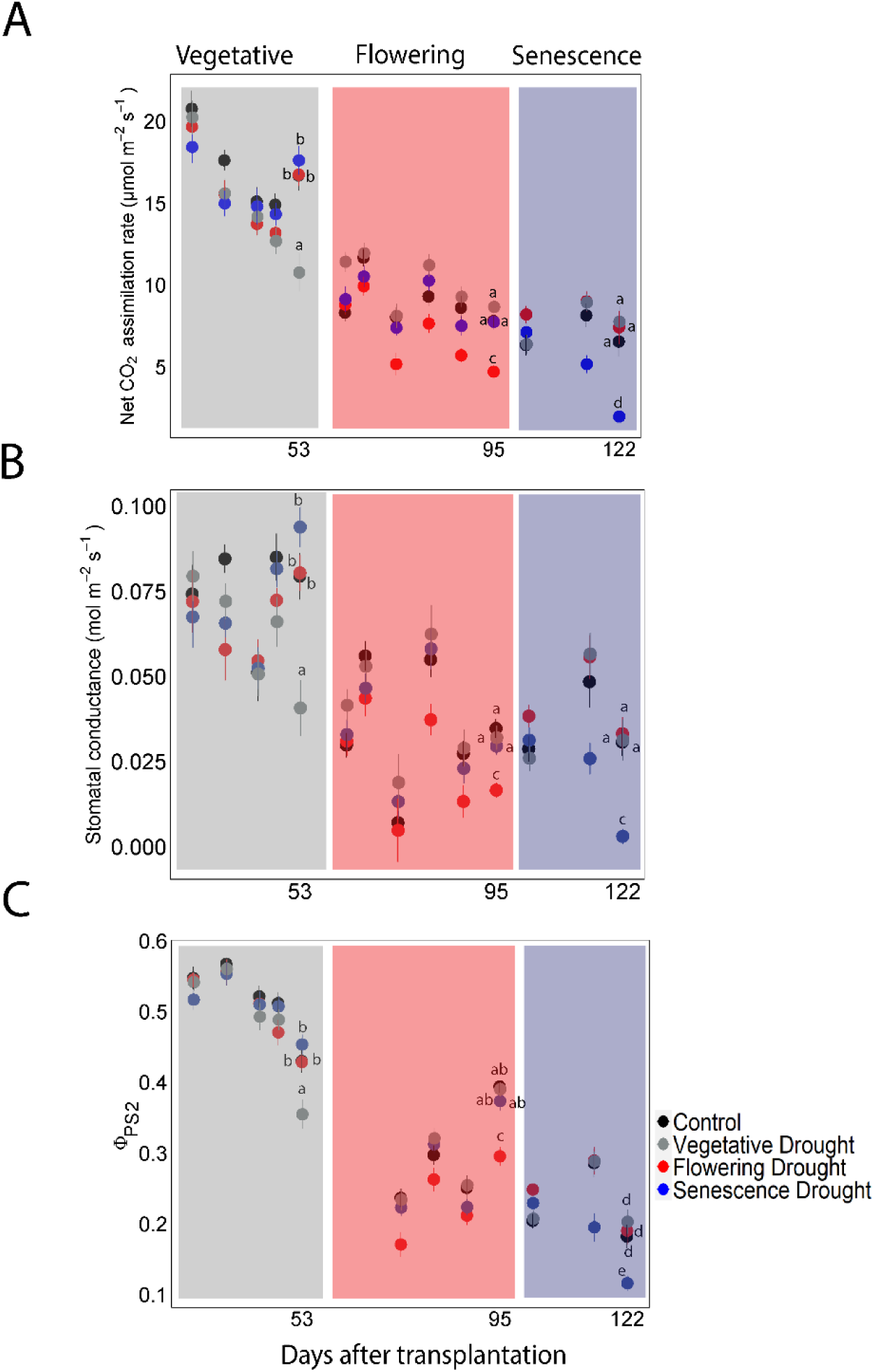
Gas exchange and Chlorophyll fluorescence from survey measurement. A. CO2 assimilation rate, B. stomatal conductance and C. operating efficiency of Photosystem II (ΦPS2) of switchgrass were measured using an LI-6800 infrared gas analyzer with fluorometer. Solid circles are averages with different colors representing the treatment when drought was implemented (N=20). Shaded regions represent developmental stages (grey = vegetative stage, red = flowering stage, blue = senescence stage). Bars on the circles represent standard error of mean. Means with different letters are significantly different according to an ANOVA and a Tukey’s post-hoc test (p<0.05)

### Switchgrass growth remained mostly robust under drought

To evaluate the impact of drought timing on switchgrass growth, key parameters including leaf number, plant height, and tiller numbers were counted weekly, with biomass yield assessed at the harvest. Drought did not affect tillering (fig. 2 A), but the number of green leaves decreased during the flowering and senescence stages (fig. 2B). Tiller production increased during the vegetative stage and ceased as the plants transitioned to the flowering stage. Biomass yields remained unaffected across treatments (fig. 2 C-E).

**Fig. 2.**
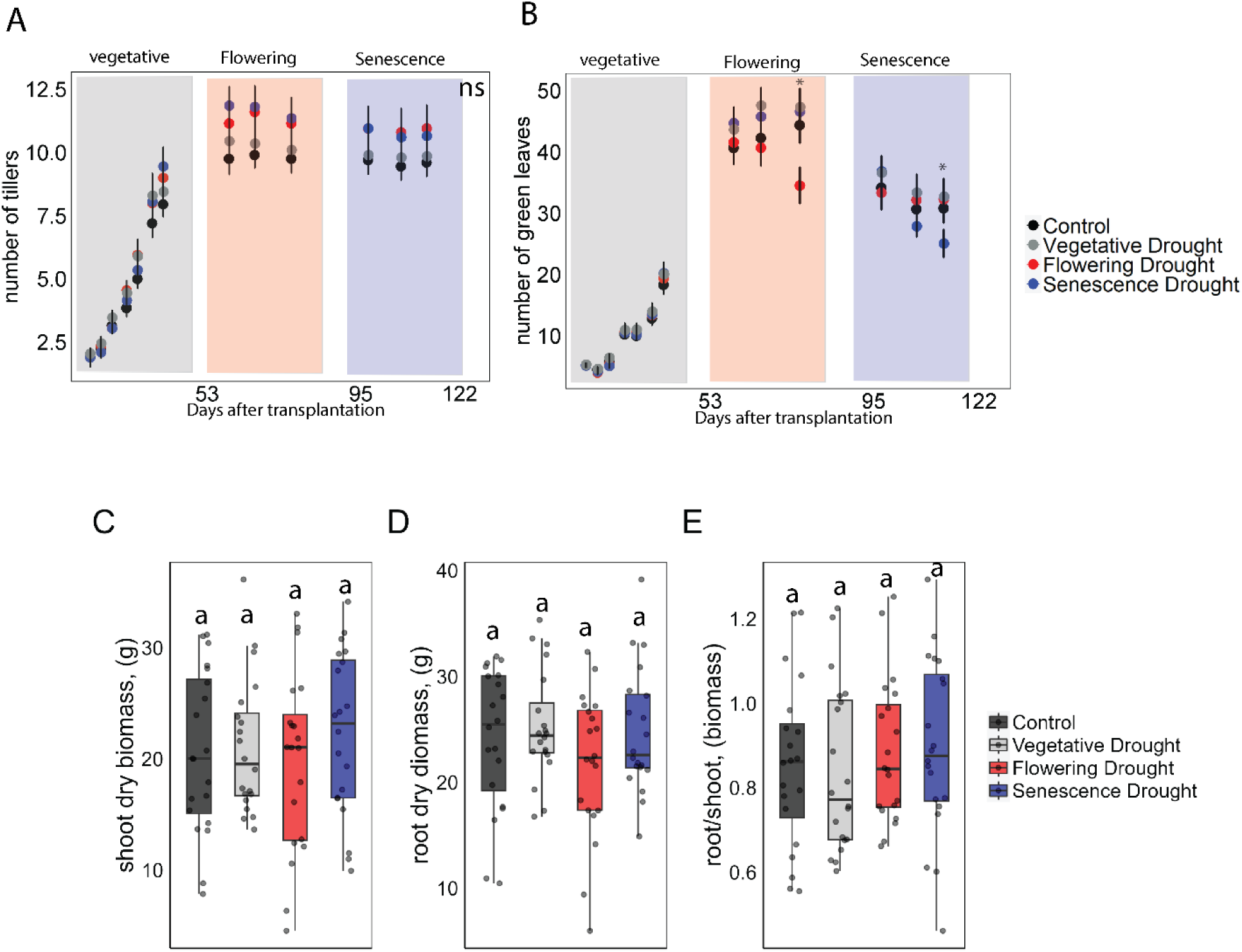
Switchgrass growth. A. Number of tillers, B. green leaves, C. shoot dry biomass, D. root dry biomass, and E. root to shoot ratio of switchgrass. Leaves and tillers were counted on a weekly basis whereas biomass was measured after the final plant harvest at the end of the experiment. Shaded regions represent developmental stages (grey = vegetative stage, red = flowering stage, blue = senescence stage). Solid circles are averages, N= 20, and vertical bars indicate standard error of mean. Asterisks indicate timepoints at which treatment groups are significantly different by Tukey’s post-hoc test (p<0.05).

### Metabolic response of switchgrass to drought across developmental stages was varied

To investigate how drought timing affects switchgrass metabolism, fresh leaf samples were collected during peak drought at each developmental stage for metabolomics analysis. This sampling included the droughted treatment, as well as all other treatments. Peak drought was defined as the point when there was a significant decline in net CO₂ assimilation.

Principal Component Analysis (PCA) of the central metabolites identified by GC-MS revealed no clear separation between treatments (fig. 3A) though several metabolites were enriched under vegetative drought treatment (fig. 3B). Differential analysis comparing control and droughted samples revealed that at the vegetative stage, levels of glucose fructose, quinnic acid, shikimic acid, and GABA increased significantly relative to controls by log2 fold-change of 2.8, 2.8, 3, 2.3, and 3.2 respectively (Fig. 4A). The effects did not persist after re-watering. At the flowering stage, central metabolite levels remained stable across treatments (fig 4 B). Interestingly, the only metabolite to show a change in abundance during senescence stage following the peak drought was succinate, which increased by a log2 fold of 2.3 (fig. 4 B,C). No single metabolite consistently responded to drought across all stages, suggesting a developmentally-specific rather than a core metabolic drought response.

**Fig 3.**
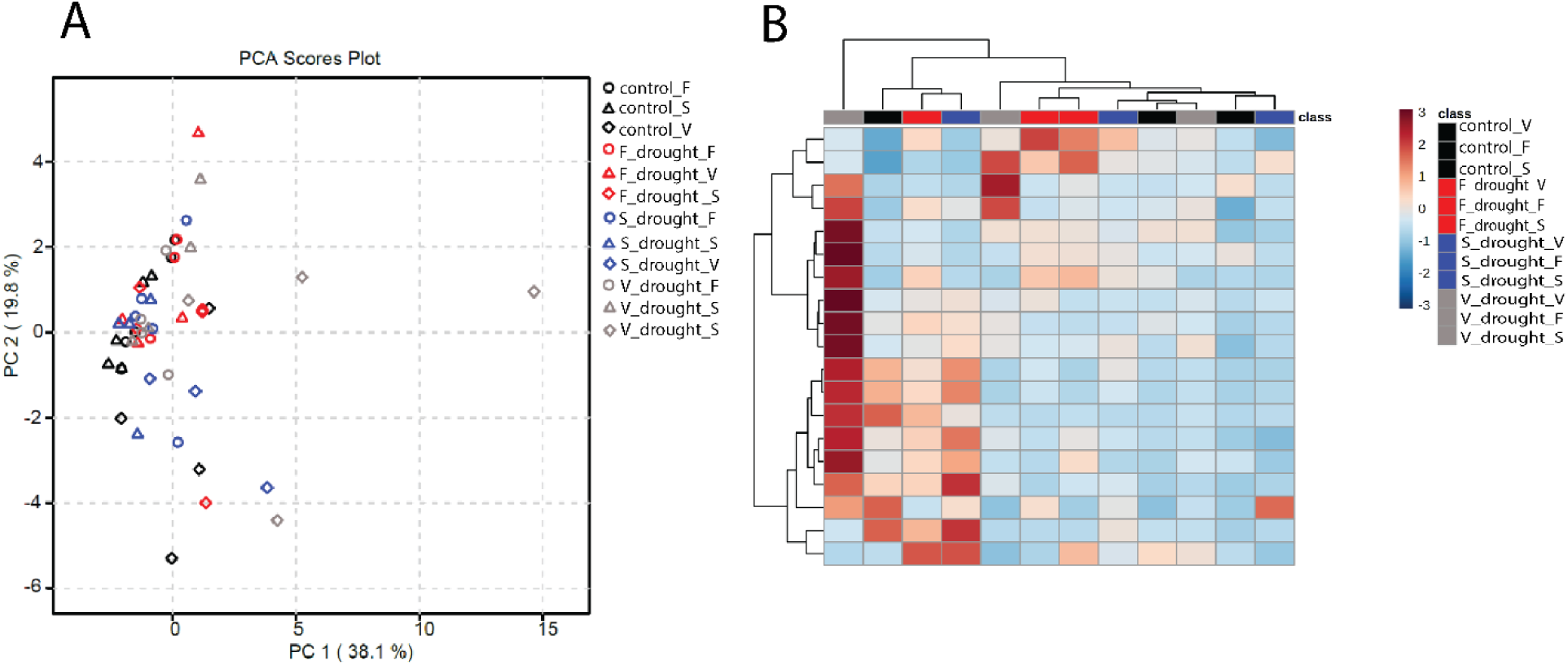
Central metabolic response to drought timing. PCA Principal Component Analysis of central metabolites identified by GC-MS A. PCA was performed on leaf secondary metabolite profiles obtained from GC-MS. The plot shows the distribution of samples along the first two principal components (PC1 and PC2), which together explain 57.9% of the total variance. Each point represents a sample, colored by treatment group, with closer points indicating greater similarity in metabolite composition. Diamond = vegetative stage, Circle = flowering stage, triangle = senescence stage. Also shown is a heatmap of leaf central metabolites identified by GC-MS B. The heatmap displays the relative abundance of central metabolites derived from switchgrass lignocellulose (N = 5) detected across different samples using GC-MS. Color intensity represents relative compound abundance, with red indicating higher concentrations and blue indicating lower concentrations. The letters _V, _F and _S at the end of the labels represent developmental stage and while that precedes these letters represent treatment. For example, F_drought_V represent that the sample received flowering drought treatment and the sampling time was at vegetative stage.

**Fig. 4.**
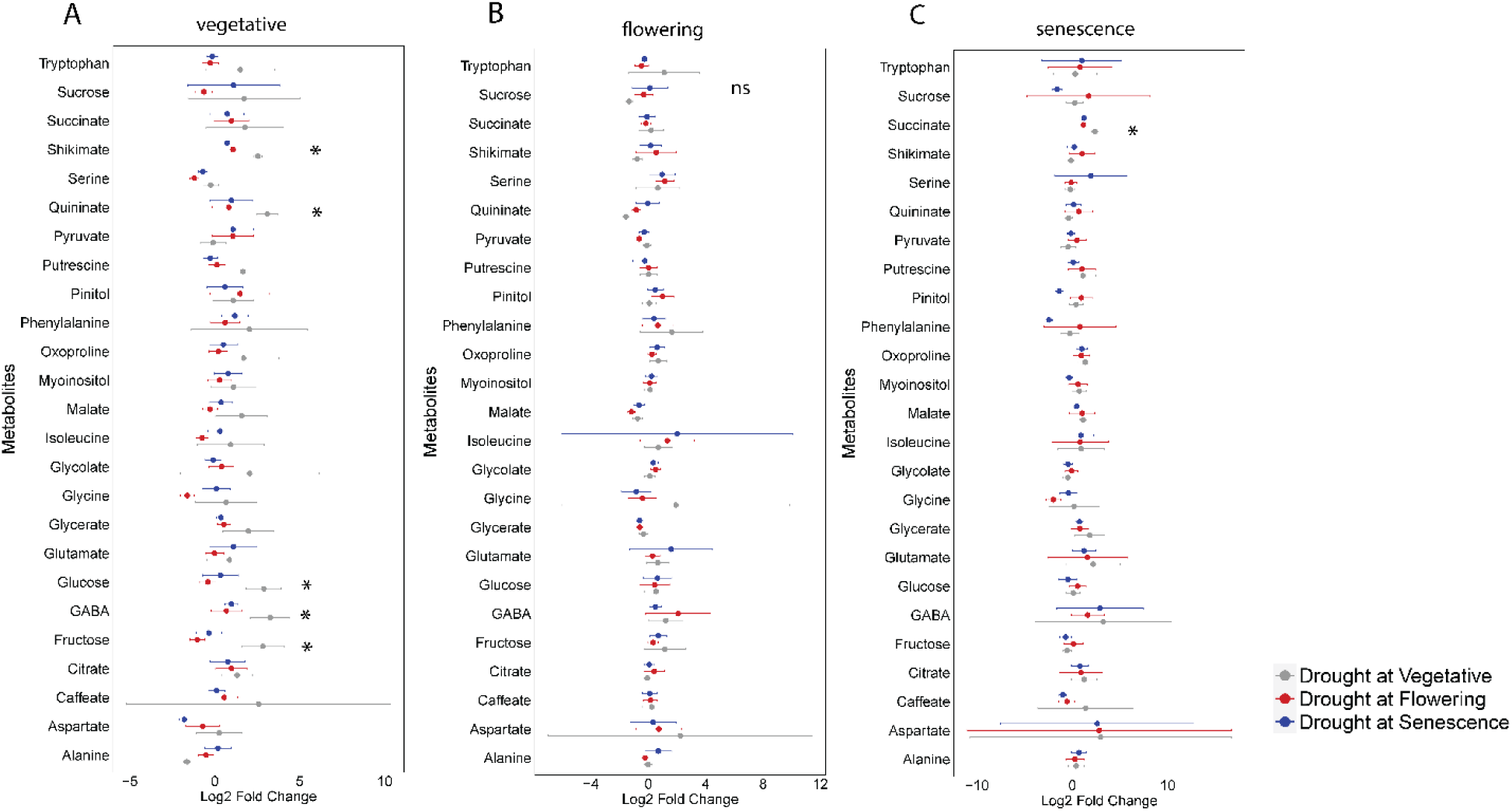
Fold change of central metabolites across three developmental stages A. Vegetative, B. Flowering, and C. Senescence. Metabolite concentrations identified using GC-MS and normalized sequentially by fresh weight and the internal standard ribitol. The data were then normalized to the control and log₂-transformed. Asterisks indicate significant differences in metabolite concentrations between drought treatments (N = 10 ±SE).

Untargeted LCMS was conducted to examine the metabolomic profiles of semi- and non-polar metabolites across treatments and developmental stages (fig 5). PC1 and PC2 together explained 32.7% (PC1-26.4%, PC2-6.3%) of the total variance. (fig. 5 A). Both PCA and hierarchical clustering showed clear separation by developmental stage, with flowering and senescence stages clustering together, and vegetative stage samples distinctly separated (Fig. 5A, B). Like the central metabolites measured using GC-MS, drought-specific metabolic shifts were only evident during the vegetative stage (Fig. 5A, C). Key metabolites driving the shift included organic acids, amides, phosphatidylcholines, and phenolics, such as flavonoids, and benzenoids, all of which responded to drought in previous work^11,12^ (fig. 5 C,D). Drought significantly enriched 235 metabolite features and depleted 427 features during the vegetative stage (Supplementary fig. 3A). Drought-induced metabolic changes were strongest at the vegetative stage, with minimal effects later, and no single metabolite consistently responded across all developmental stages.

**Fig. 5.**
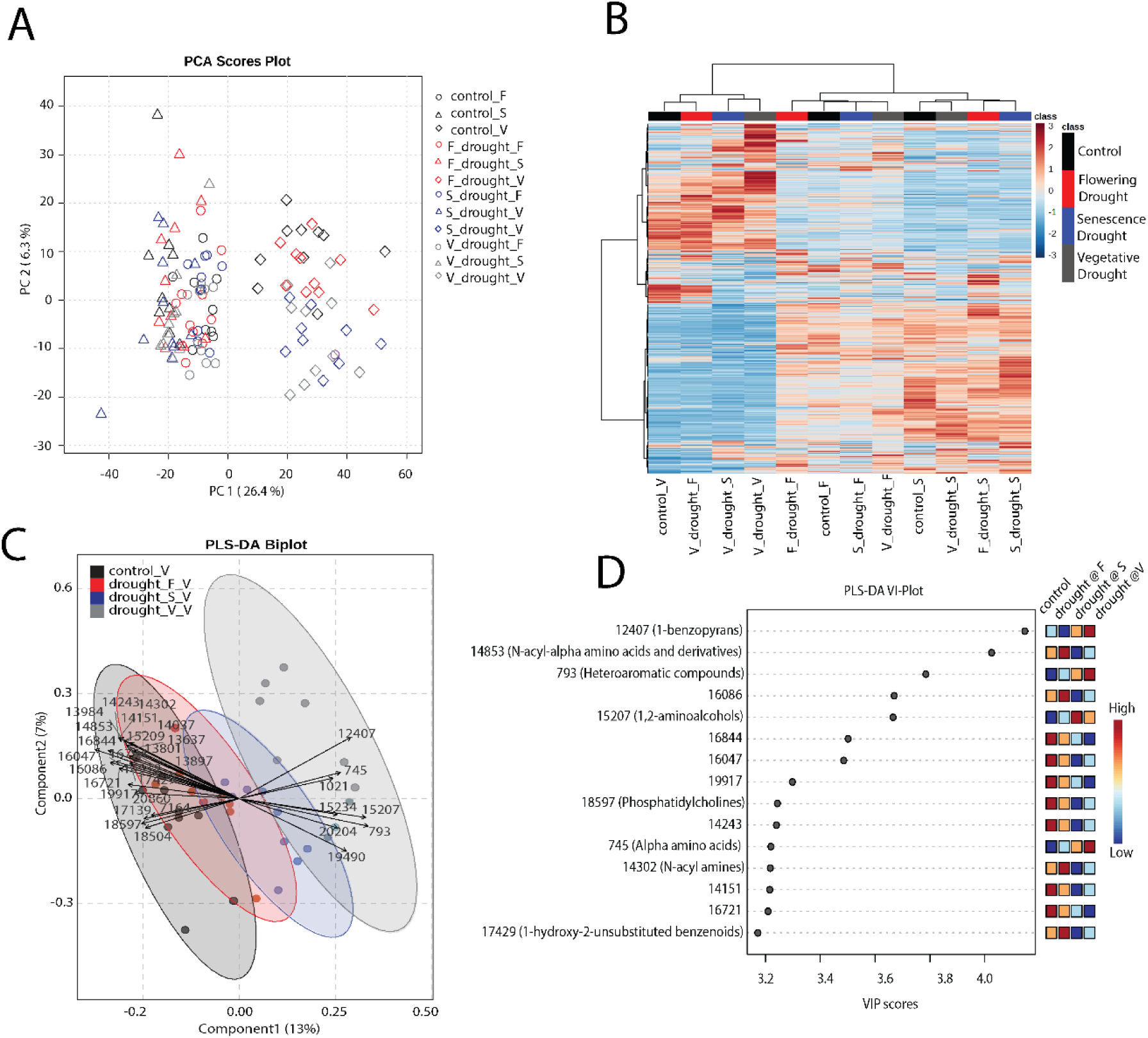
Specialized metabolic response to drought timing A. PCA Principal Component Analysis of secondary metabolites identified by LC-MS. PCA was performed on leaf secondary metabolite profiles obtained from LC-MS. The plot shows the distribution of samples along the first two principal components (PC1 and PC2), which together explain 32.7% of the total variance. Each point represents a sample, colored by treatment group, with closer points indicating greater similarity in metabolite composition. B. Heatmap of hydrolysate compounds identified by LC-MS. The heatmap displays the relative abundance of secondary metabolites derived from switchgrass leaves (N = 10) detected across different samples using LC-MS. Color intensity represents relative compound abundance, with red indicating higher concentrations and blue indicating lower concentrations. The letters _V, _F and _S at the end of the labels represent developmental stage and while that precedes these letters represent treatment. For example, F_drought_V represent that the sample received flowering drought treatment and the sampling time was at vegetative stage. Diamond = vegetative stage, Circle = flowering stage, triangle = senescence stage. Each cell on the heatmap represents an averaged metabolite abundance from 10 (biologically) replicated samples. C. Partial Least Squares Discriminant Analysis for the samples collected at the vegetative stage. The two components explained 20% of the total variance. D. Variable Importance Plot showing the most important metabolites that drove the metabolite profile separation.

### Senescence drought enhanced ethanol production from hydrolysis

Previous studies indicated that yeast fermentation of hydrolysates from drought-stressed switchgrass was severely impaired^6,8^. To determine how drought timing impacts yeast fermentation and whether the effects persist across developmental stages, switchgrass samples were subjected to Soaking in Aqueous Ammonia (SAA) pretreatment and enzymatic hydrolysis. Hydrolysates were then fermented using engineered *S. cerevisiae*. Interestingly, ethanol production was highest in senescence drought treatment with 62% increase compared to the control and 95% and 87% increases compared to the vegetative and flowering drought groups (fig. 6A). Consistently, glucose consumption by yeast remained highest in the senescence drought group (56% higher compared to control and 89% and 82 % higher compared to vegetative senescence drought groups respectively, fig. 6B). Cell growth (OD_600_) was significantly inhibited by vegetative (97 % lower) and flowering drought (83% lower) treatments (fig. 6C, Supplementary fig. 4). Similarly, a delay in fermentation occurred in vegetative stage compared to senescence stage, although, there was no difference as compared to control or plants droughted during the flowering stage (supplementary fig. 5)

**Fig. 6.**
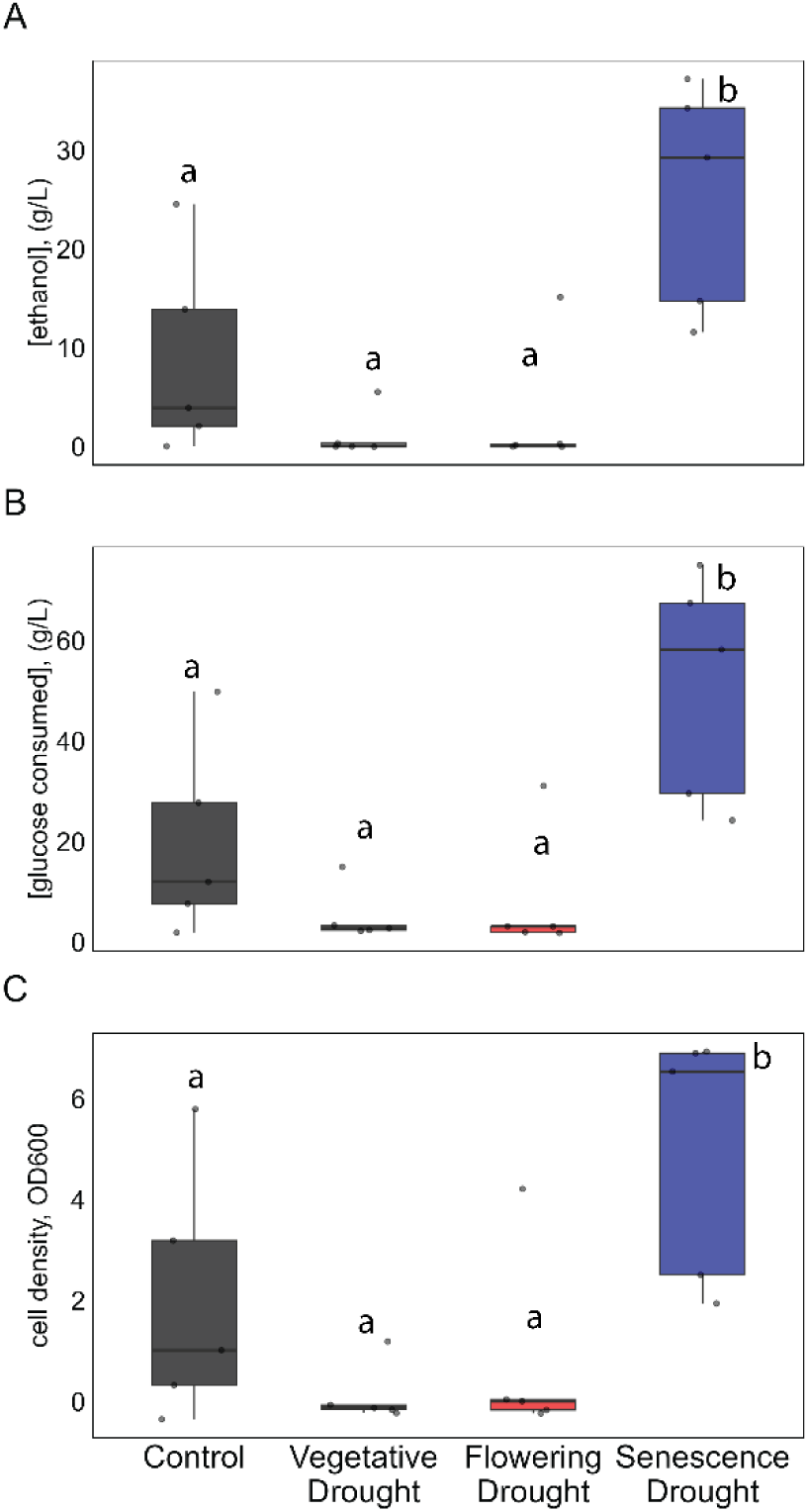
Fermentation results by yeast with hydrolyzed switchgrass subjected to drought at different developmental stages. A. Ethanol titers from senesced aboveground biomass of switchgrass using *Saccharomyces cerevisiae*. B. Cell density of *S. cerevisiae* and C. amount of glucose consumed during fermentation of hydrolysates. Bars show averages, N= 5, and vertical bars indicate standard error of mean. Groups with dissimilar letters are significantly different by Tukey’s post-hoc test (p<0.05).

To determine causes of inhibition, we analyzed the hydrolysates, focusing first on the previously known lignocellulose-derived inhibitors. A heatmap revealed differential enrichment patterns across the treatments (fig. 7) with 12 compounds showing significant difference between treatments (fig. 8). In senescence drought treatment, 3,4-dihydroxybenzaldehyde, azelaic acid, benzamide, delta valerolactum, ferulic acid, feruloyl amide, kynurenic acid, phthalic acid, and suberic acid exhibit significantly lower levels compared to at least one other treatment, while 4-hydroxybenzaldehyde, N-methylnicotinamide, and xanthurenic acid show either elevated or similar levels compared to the other treatments.

**Fig. 7.**
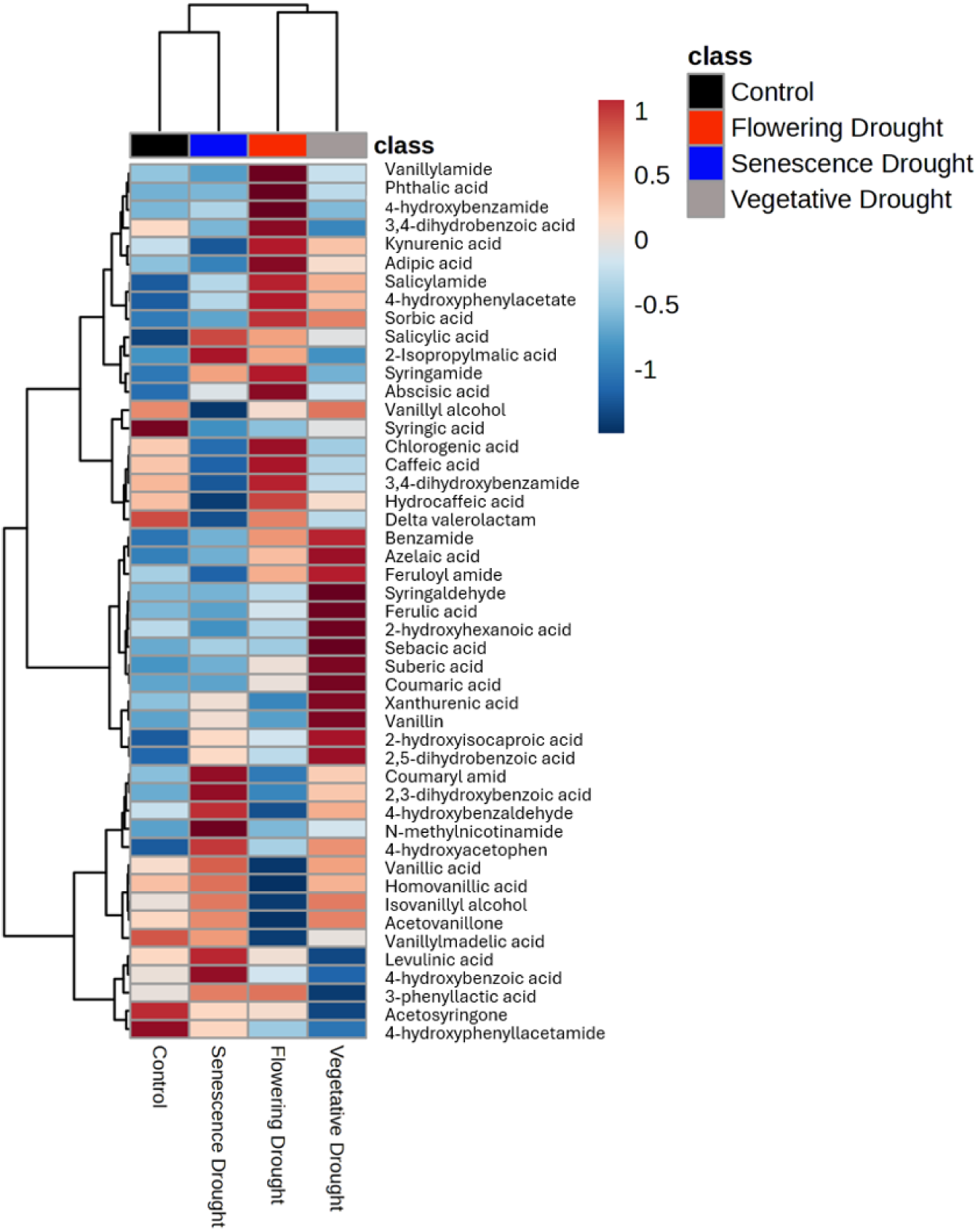
Heatmap of hydrolysate compounds identified by LC-MS. The heatmap displays the relative abundance of hydrolysate compounds derived from lignocellulose and detected across different samples using LC-MS. Compounds were identified (N = 5) and quantified based on their retention time and m/z values, followed by data normalization and hierarchical clustering to reveal patterns of variation among samples. Color intensity represents relative compound abundance, with red indicating higher concentrations and blue indicating lower concentrations.

**Fig. 8.**
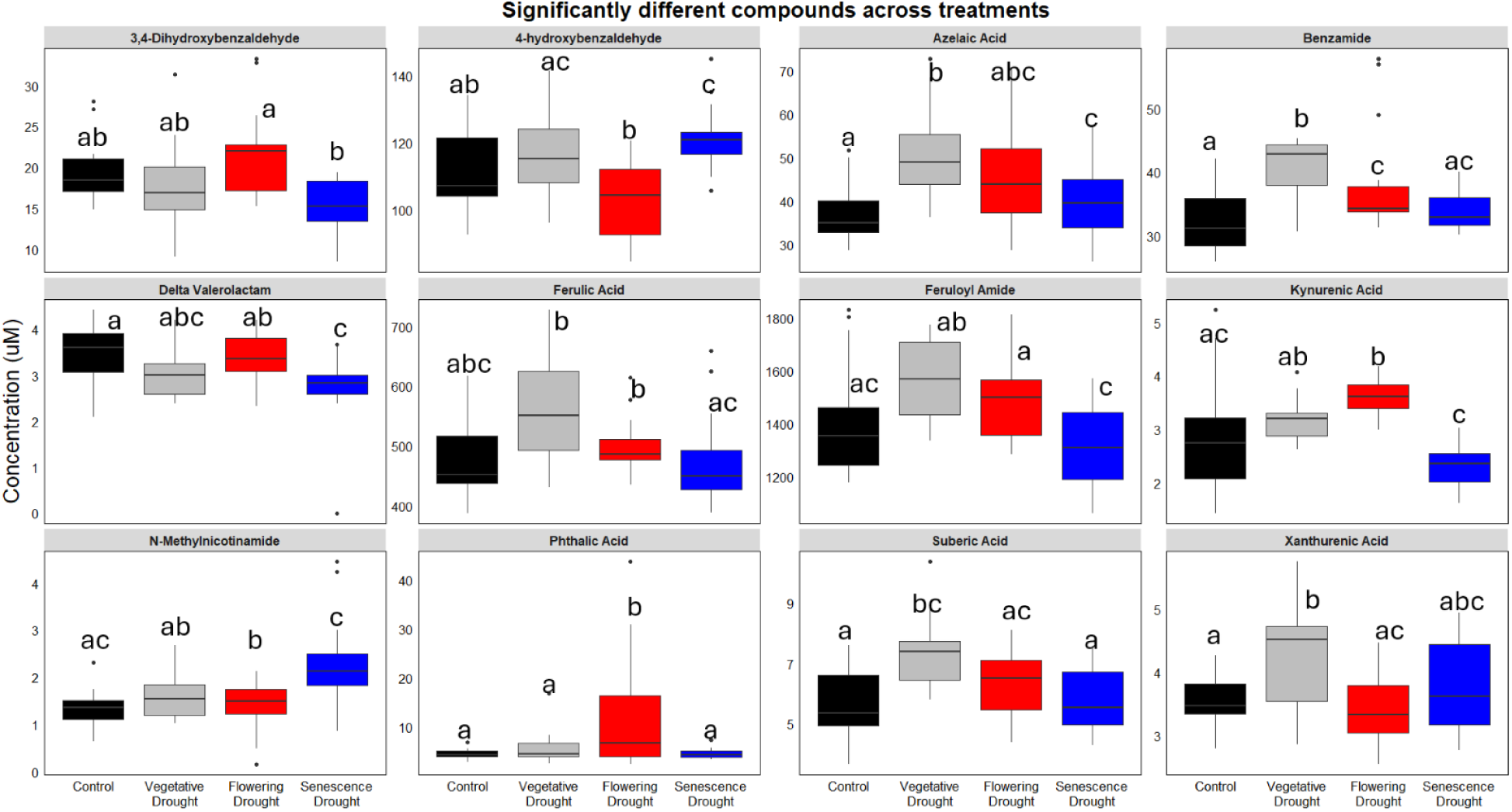
Concentration of switchgrass hydrolysate compounds that are significantly different among treatment groups. The boxplot displays the distribution of concentration of hydrolysate compounds derived from lignocellulose and detected across different samples using LC-MS. Compounds were identified (N = 5) and quantified based on their retention time and m/z values. The boxes show the interquartile range (IQR, the middle 50% of values), the horizontal line inside the box represents the median, and the whiskers indicate the data spread. Outliers are displayed as individual points.

Given recent evidence^8^ suggesting that drought-induced saponins may inhibit yeast growth during fermentation, we conducted untargeted metabolomics to assess saponin presence in the hydrolysates. We identified five unique saponins in the hydrolysates out of approximately 60 originally identified in fresh biomass. Notably, the total saponin level was lowest in senescence drought hydrolysates (fig 9 A) showing a 36% reduction compared to the control and 54% and 59% lower levels compared to vegetative and flowering drought treatments. In contrast, drought during earlier stages elevated total saponin levels, although the increases were not statistically significant. Mixed effect modeling revealed that hydrolysate saponin levels strongly predicted fermentation lag time with R^2^ values ranging from 0.39 to 0.59 (fig. 9B). Prolonged lag phases (defined as time the function takes to reach 10% of its maximum) observed with the high saponin-containing samples suggest that they may inhibit yeasts acclimating to the hydrolysate environment (supplementary fig. 5).

**Figure 9.**
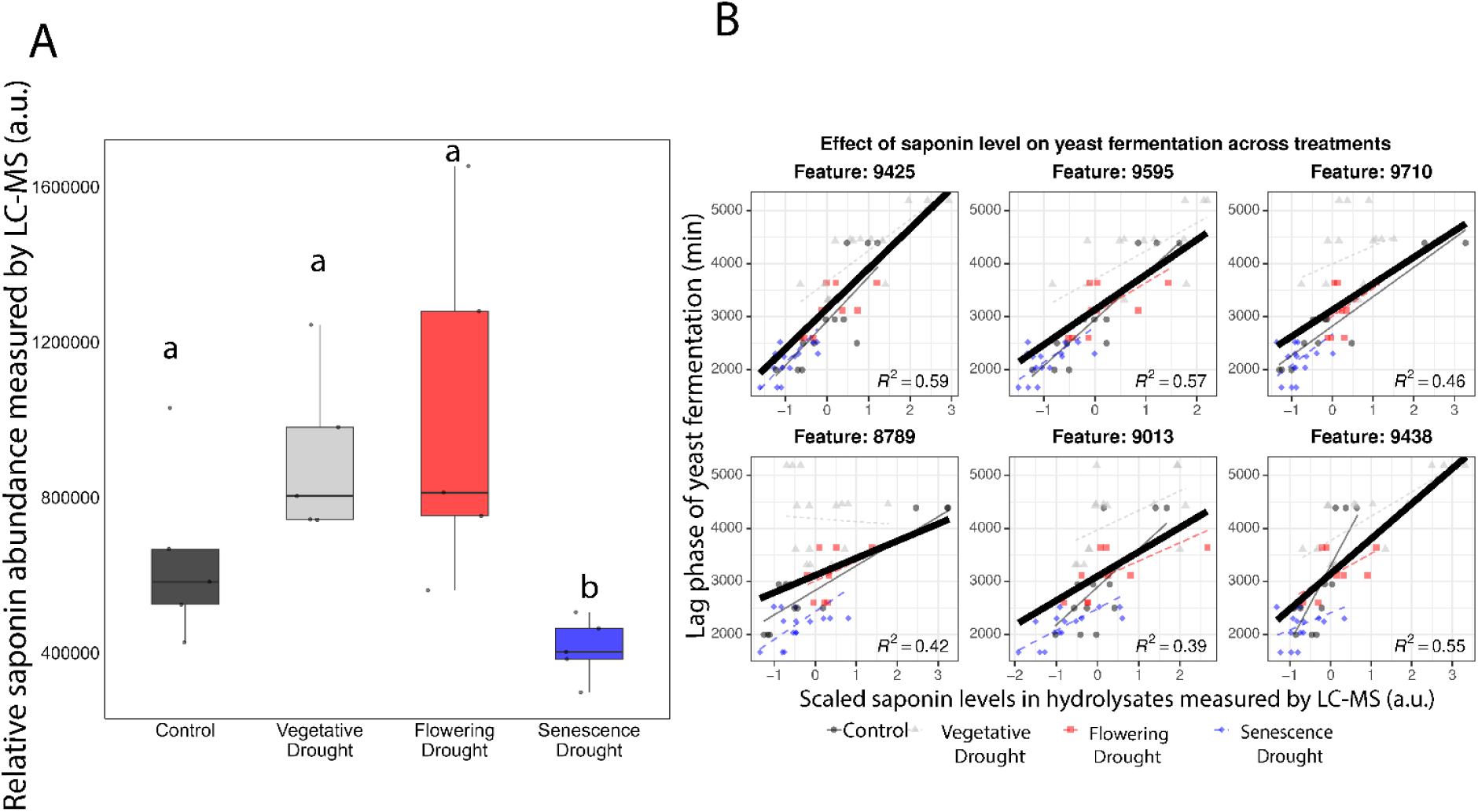
A. Relative abundance of total saponins found in hydrolysates. Bars show averages, N= 5, and vertical bars indicate standard error of mean. Data was normalized to internal standard telmisartan relative abundance. Groups with dissimilar letters are significantly different by Tukey’s post-hoc test (p<0.05). B Scatter plot of lag time as a function of scaled saponin content, with linear regression lines shown per treatment group. Lines reflect random slopes from a mixed-effects model (*Lag ∼ Mass_scaled + (Mass_scaled | treatment_group)*)

In addition to saponins, untargeted metabolomics identified 91 unique metabolite features that positively correlated with the lag phase time (>0.5 correlation, FDR p ≤ 0.05, supplementary table 1), suggesting their potential inhibitory roles to the yeasts. Chemical annotation indicated that these features were chemically diverse including terpenoids, steroids, phenolics, alkaloids, amino acids (and derivatives) and fatty acid derived metabolites (supplementary fig. 6, supplementary table 1). Nitrogen-containing compounds, like amino acids (and derivatives), different types of alkaloids, amides and amines, were overrepresented among the annotated features. Among all compounds, saponins correlated most strongly to fermentation lag times, highlighting their central role in drought-related inhibition of yeast fermentation.

## Discussion

This work demonstrates that switchgrass (*Panicum virgatum* L.) grows robustly under drought, despite significant reduction of photosynthetic parameters and regardless of when the drought is imposed developmentally. Reduced stomatal conductance under drought stress emerged as a principal factor likely diminishing CO₂ assimilation (fig. 1 A-B). During severe drought, stomatal closure is an initial mechanism to minimize water loss, but this concurrently restricts CO₂ entry into intercellular spaces and bundle sheath cells, leading to decreased photosynthetic rates, even in C4 plants ^13^. In this study in switchgrass, these detrimental effects become pronounced only under extreme soil dryness, when soil moisture dropped to 1% VWC for extended periods of time (supplementary fig. 1). At this soil moisture, a significant reduction in leaf water potential occurred (supplementary fig. 2). This observation underscores switchgrass tolerance to extreme drought conditions, where clear physiological responses are typically elicited only when soil moisture is critically low or when drought conditions are prolonged. In addition to the drought effects, a progressive decline in net CO_2_ assimilation and Φ_PS2_ is also observed over time (fig. 1A and C). This decline in photosynthesis can be explained by potential carbon sink limitation as the plant matures. When vegetative growth is completed, starch reserves in rhizomes are filled which diminishes rhizome sink activity, resulting in low carbon demand from photosynthesis ^10^. There was also a significant depletion of leaf starch at the senescence stage, potentially due to declining photosynthesis and the plant’s requirement of starch as energy reserve for cellular maintenance (supplementary fig. 7). These photosynthetic responses comprise a core physiological response to drought.

Despite the reduction in photosynthetic capacity under severe drought stress, there were no significant differences in either aboveground and belowground biomass yield or tillering (fig. 2 A, C, and D). The only growth response to drought was a reduced number of green leaves when implemented at the flowering and vegetative stage (fig. 2B). This resilience may be attributed to the experimental design wherein plants subjected to extreme drought were promptly rehydrated to ambient soil moisture levels and maintained under these conditions for the remainder of the experimental period. The short duration of the drought treatment, albeit intense, likely allowed the plants to recover and maintain normal growth processes ^14^. This suggests that switchgrass possesses effective mechanisms to withstand short-term severe drought without compromising overall biomass production. Stomatal conductance, on the one hand, is the first plant response to depleting soil moisture and can thus concurrently reduce CO_2_ assimilation which explains why we observed dynamic response of photosynthesis to changing soil moisture ^15^. Switchgrass is highly resilient to short-term moderate to extreme moisture stress, with drought having little to no impact on yield^16,17^. However, short-term drought may interact with other environmental factors, such as soil fertility, potentially influencing biomass quality^17^. On the other hand, long-term drought over a single season or spanning a few years generally reduces aboveground biomass production in switchgrass ^18,19^ We next investigated the potential metabolic underpinnings to this high drought resilience.

In central metabolism, switchgrass responds somewhat to drought during two developmental stages (fig. 3 A,B). Drought during the vegetative stage elicited the most pronounced changes, followed by the senescence stage which is marked only by a single metabolite response to drought (4 A,C). Interestingly, the flowering stage was least responsive to drought (fig. 4 B). Under drought conditions during the vegetative stage, there was a significant accumulation of monosaccharides such as glucose and fructose, organic acids like quinate and shikimate, the non-proteinogenic amino acid gamma-aminobutyric acid (GABA), and the dicarboxylic acid succinate (fig. 4 A). These metabolites play crucial roles in stress adaptation; for instance, glucose and fructose function as osmolytes, helping to maintain cell turgor and prevent dehydration ^20^. Additionally, with photosynthesis being compromised, the plant may increasingly rely on stored sugars as alternative energy sources ^21^, hence, necessitating the accumulation of sugars. For an example, in soybean leaves at vegetative stage, drought causes an increase in accumulation of soluble sugars with a concomitant decrease in starch ^22^. GABA is a key player in drought resistance in plants. Accumulation of GABA enhances drought tolerance mainly by regulating stomatal opening, inhibiting reactive oxygen species production, activating antioxidation and enhancing photosynthesis ^23^. It is integral to the GABA shunt pathway, which aids in integrating carbon and nitrogen metabolism, thereby supporting energy production and essential metabolic processes when normal photosynthesis is impaired ^24^. Similarly, shikimate is a precursor for the production of phenolics, some of which are associated with drought tolerance ^24^. Among organic acids, succinates are one of the most responsive metabolites to drought ^25^. Succinic acid, or succinate, serves as a key intermediate in the ATP-generating Tricarboxylic Acid Cycle (TCA), which is essential for energy production and mitochondrial regulation. Under drought stress, elevated NADH levels inhibit several TCA cycle dehydrogenases (pyruvate dehydrogenase, isocitrate dehydrogenase, α-ketoglutarate dehydrogenase, and citrate synthase), while succinate dehydrogenase remains unaffected, converting succinyl-CoA to succinate. This overproduction of succinate in our droughted samples potentially enables mitochondrial ATP accumulation to sustain cellular functions under drought conditions from stored sugars ^26^.

Our results revealed much larger and more extensive changes in specialized metabolite profiles between developmental stages as compared to central metabolism, but still mainly during the vegetative stage drought. During the vegetative phase, metabolite differences associated with drought stress were also clear (fig. 5 B and supplementary fig. 3). The most important metabolites that drove the developmental stage specific shift include 1-benzopyrans, N-acyl-alpha amino acids and derivatives, 1,2-aminoalcohols, phosphatidylcholines, alpha amino acids, N-acyl amines and 1-hydroxy-2-unsubstitued benzenoids (fig. 5 C, D). Most of these metabolites have important biological roles in plants such as plant metabolism, defense and stress responses^27–29^.Comparing specialized metabolite profiles between control and drought-treated groups however, revealed no significant metabolic response during the flowering or senescence stages (supplementary fig. 3 B, C). Moreover, increasing the false discovery rate threshold from 5% to 10% did not result in any features being significantly different at these two stages. The potential explanation could be that the vegetative stage was characterized by rapid growth and high metabolic activity, enabling strong adaptive responses to drought, including changes in gene expression, protein synthesis, and enzyme activity that enhance drought tolerance. In contrast, during the flowering and senescence stages, photosynthetic activity decreased as resources are redirected toward reproductive development and the completion of the life cycle of switchgrass for the season. Consequently, the metabolic response to drought stress was less pronounced during these stages, as the plant may have prioritized reproduction over stress tolerance.

Although the physiological responses to the various drought timings were similar, the metabolic responses were not. In this study, there was no single, core metabolic pattern across the treatments, indicating that different biochemical processes were at play depending on the timing of the drought. Drought during senescence increased fermentation relative to both control and other drought timing treatments. (fig. 6). This was evidenced by reduced yeast cell density and lower CO₂ production rates, indicating reduced fermentation in these samples compared to those droughted during senescence (fig. 6 B and C). The higher concentrations of saponins in the hydrolysates from control, vegetative, and flowering drought samples may have contributed to this inhibition (fig. 9 A), given the antifungal properties of switchgrass saponins against switchgrass root associating fungi and their ability to impede yeast fermentation ^8^. This observation is further supported by the mixed model analysis, which shows that saponin levels in the hydrolysates strongly predict the fermentation time lag, as indicated by a strong R-squared value (fig. 9 B). Switchgrass exposed to drought accumulates saponins^8^ and fermentation of biomass from this drought-stressed switchgrass using *Saccharomyces cerevisiae* yields significantly lower ethanol compared to that from biomass obtained under ambient conditions^6^. Moreover, saponins added exogenously to the hydrolysates obtained from switchgrass grown under normal rainfall suppressed the fermentation activity of *S. cerevisiae*, confirming the inhibitory effects of saponins on ethanol production^8^. Our study is the first to exhibit a robust link between fermentation inhibition and drought induced saponins across a variety of treatments.

Similar to the leaf level metabolites, there was a developmentally-specific drought response in hydrolysate compounds. The variety of lignocellulose-derived inhibitory compounds that were identified in the hydrolysates showed significant accumulation in response to drought during the vegetative and flowering stages (fig. 7). In total, 12 out of 50 hydrolysate compounds in the hydrolysates showed significant differences between treatments (table 1. fig. 8). Feruloyl amide and coumaroyl amide, which form when ferulic acid and *p*-coumaric acid react with ammonia during the ammonia pretreatment, were present in disproportionately larger amounts compared to other compounds (Supplementary fig. 5). These two compounds are known fermentation inhibitors ^9,30^. Across treatments, the concentration of coumaroyl amide was similar whereas feruloyl amide was significantly higher in Vegetative drought group and lower in Senescence drought group (supplementary fig. 8). A certain concentration of inhibitors is necessary for toxicity to occur. It is possible that coumaroyl amide, at the levels found in our samples, is not problematic on its own. However, the combined presence of coumaroyl amide, feruloyl amide, saponins, or other compounds may work together to synergistically inhibit fermentation. The Senescence drought group may have a more favorable combination of the toxic compounds, which could explain why, despite containing many inhibitory compounds, this group showed the least inhibition of fermentation. Notably, among the potential inhibitors measured in this study, saponins showed the strongest effect on fermentation inhibition, as evidenced by their ability to most strongly predict fermentation lag time (fig. 9 B). This suggests that saponins, more than other hydrolysate compounds, played a dominant role in fermentation suppression. The Senescence drought group, which exhibited the lowest saponin levels, also experienced the least fermentation inhibition, further reinforcing the idea that saponins were the primary driver of this effect. While other known inhibitors were present, their influence appeared secondary to the impact of saponins. These findings highlight the critical role of saponins in fermentation outcomes and suggest that their depletion in the Senescence drought group contributed to its more favorable fermentation outcome.

The hydrolysates predominantly included long-chain dicarboxylic acids (pimelic, sebacic, and suberic acid) along with ferulic acid and vanillin. Vanillin can inhibit microbial growth, including *Saccharomyces cerevisiae* ^31^. Ferulic acid and *p*-coumaric acid, which play a vital role in lignin structure, also accumulated more under drought conditions. In addition to reacting with ammonia to form feruloyl amide and coumaroyl amides, which inhibit fermentation, these compounds are themselves known to inhibit *S. cerevisiae* fermentation ^5,32^. Most of the amides found in the hydrolysates were potentially formed due to the reaction between organic acids and ammonia during the pretreatment process. Most of the lignocellulose-derived phenolic inhibitors in our hydrolysates are non-metabolizable by fermentative organisms. Hydrolytic treatments often exacerbate the degradation of lignin and monomeric sugars, leading to the formation of three major groups of fermentation inhibitors, furan derivatives, weak acids, and phenolic compounds ^33^. The accumulation of these compounds can substantially impede the fermentation process^34^. Our data revealed that these compounds accumulated under drought conditions across different developmental stages. Notably, no significant changes were observed in the total content or composition of cell wall polysaccharides and lignin (supplementary fig. 9-11).

These findings suggest that drought stress, particularly during the vegetative and flowering stages, induces the accumulation of specific metabolites that can inhibit fermentation processes, potentially impacting biofuel production efficiency from switchgrass biomass. The production of antifungal compounds like saponins may be part of the plant defense strategy under stress but pose challenges for industrial fermentation applications. Understanding the metabolic shifts in switchgrass under varying drought conditions and developmental stages is crucial for optimizing biomass utilization and improving biofuel production processes.

## Acknowledgments

This material is based upon work supported by the Great Lakes Bioenergy Research Center, U.S. Department of Energy, Office of Science, Biological and Environmental Research Program under Award Number DE-SC0018409. This work was also supported by the Chemical Sciences, Geoscience and Biosciences Division, Office of Basic Energy Sciences, Office of Science, U.S. Department of Energy (DE-FG02-91ER20021). We thank Dan Xie, Kallysa Taylor and Jose Serate for pretreatment, hydrolysis and fermentation services, Kurt Creamer and Novozymes for generously providing enzymes, the GLBRC Bioanalytics Facility for glucan content quantification, and the GLBRC Metabolomics Facility for quantification of analytes.

## Author Contribution

BB, BJW: conceptualization; XL, KAO: LC-MS; VJP: data collection; XF,BB: GC-MS; YZ, TKS: Fermentation; JJC, RLL: review and editing, BJW: funding acquisition

## Declarations of Interest

The genetic modifications in GLBRCY1455 are covered by patents held by the Wisconsin Alumni Research Foundation. TKS is an inventor in those patents.

## Funding Statement

This work was supported in part by the Great Lakes Bioenergy Research Center, U.S. Department of Energy, Office of Science, Office of Biological and Environmental Research under Award Number DE-SC0018409, and Basic Energy Sciences under Award DE-FG02-91ER20021.

## Data Availability

All data supporting the findings of this study are available either within the paper and its supplementary data or upon request from the corresponding author.

---

**Supplementary fig. 1.**
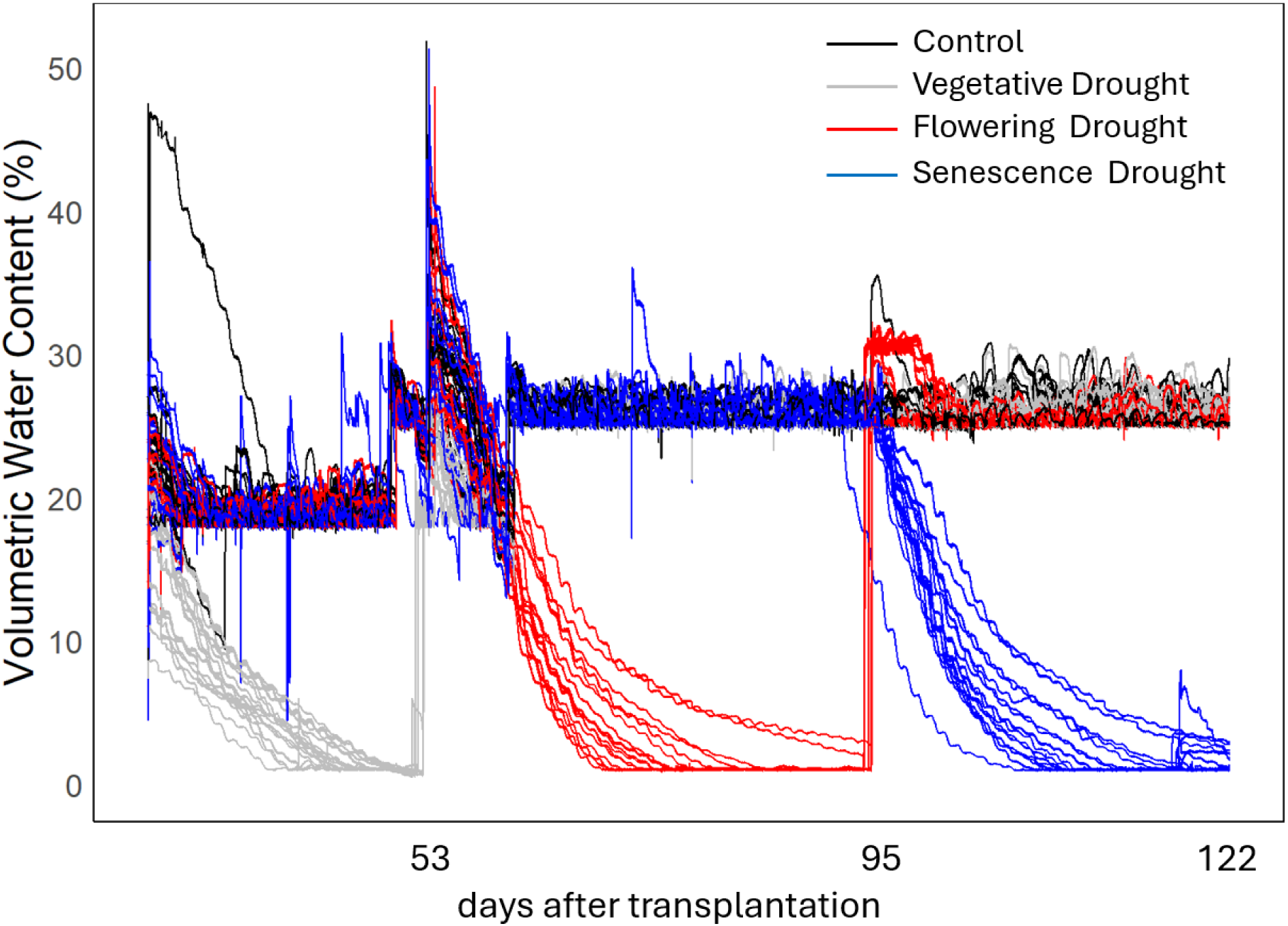
Soil moisture trends throughout the experimental period, as measured by different soil moisture sensors. Irrigation was controlled using a custom-built automated system designed to maintain soil moisture at a specific set point. Soil moisture data were collected by TEROS-12 probes, which communicated with a Raspberry Pi microcontroller via the SDI-12 interface. The Raspberry Pi triggered or halted irrigation based on whether the soil moisture fell below or exceeded the set threshold.

**Supplementary fig. 2.**
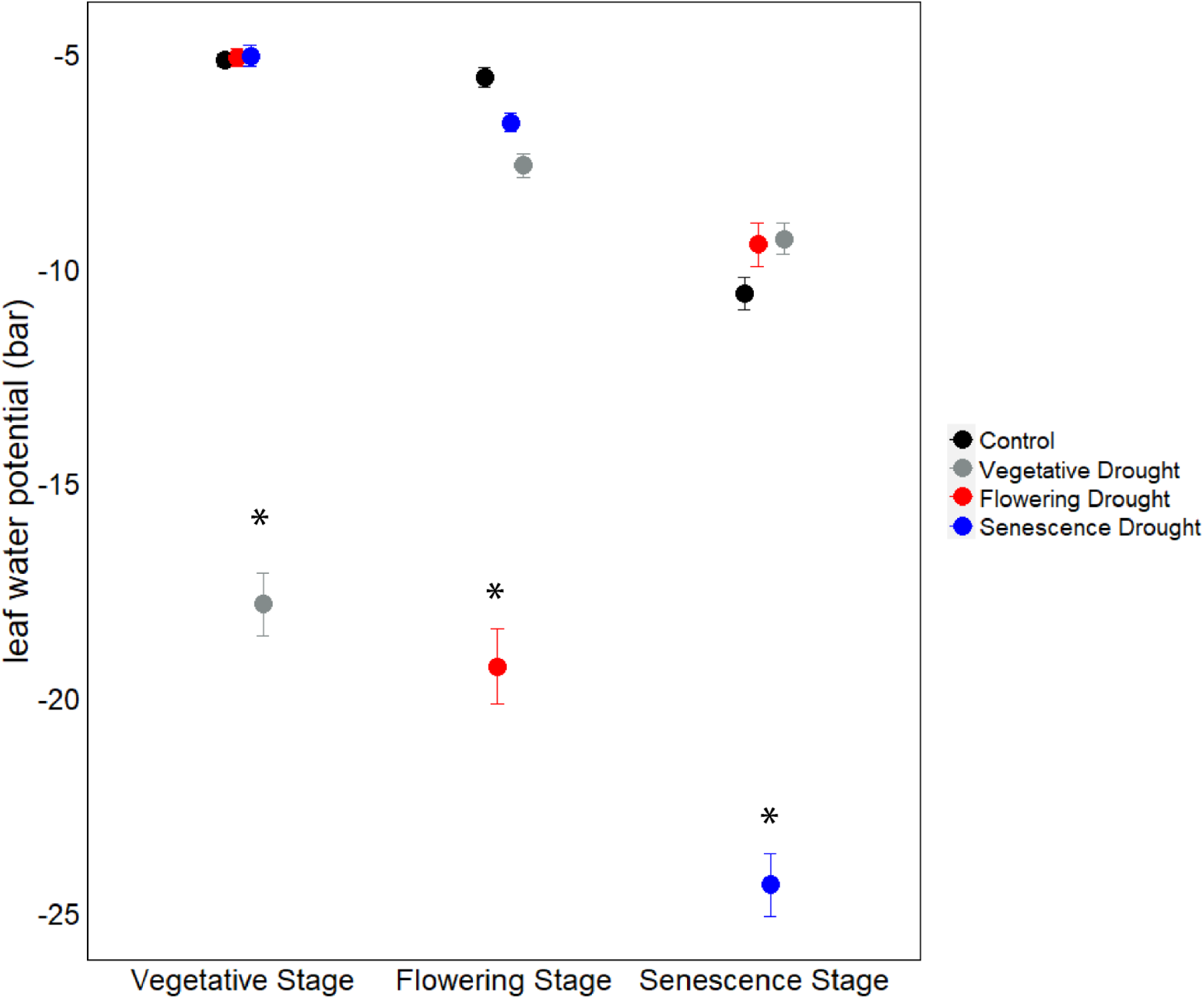
Leaf water potential of switchgrass measured during the peak drought using Scholander pressure bomb. It measures leaf water potential by applying compressed gas to a leaf’s cut stem until xylem sap emerges, indicating the pressure needed to balance the tension within the plant. Colored points represent mean values of 20 replicates each with vertical bars showing standard error of the mean. Asterisks indicate timepoints at which treatment groups are significantly different by Tukey’s post-hoc test (p<0.05).

**Supplementary fig. 3.**
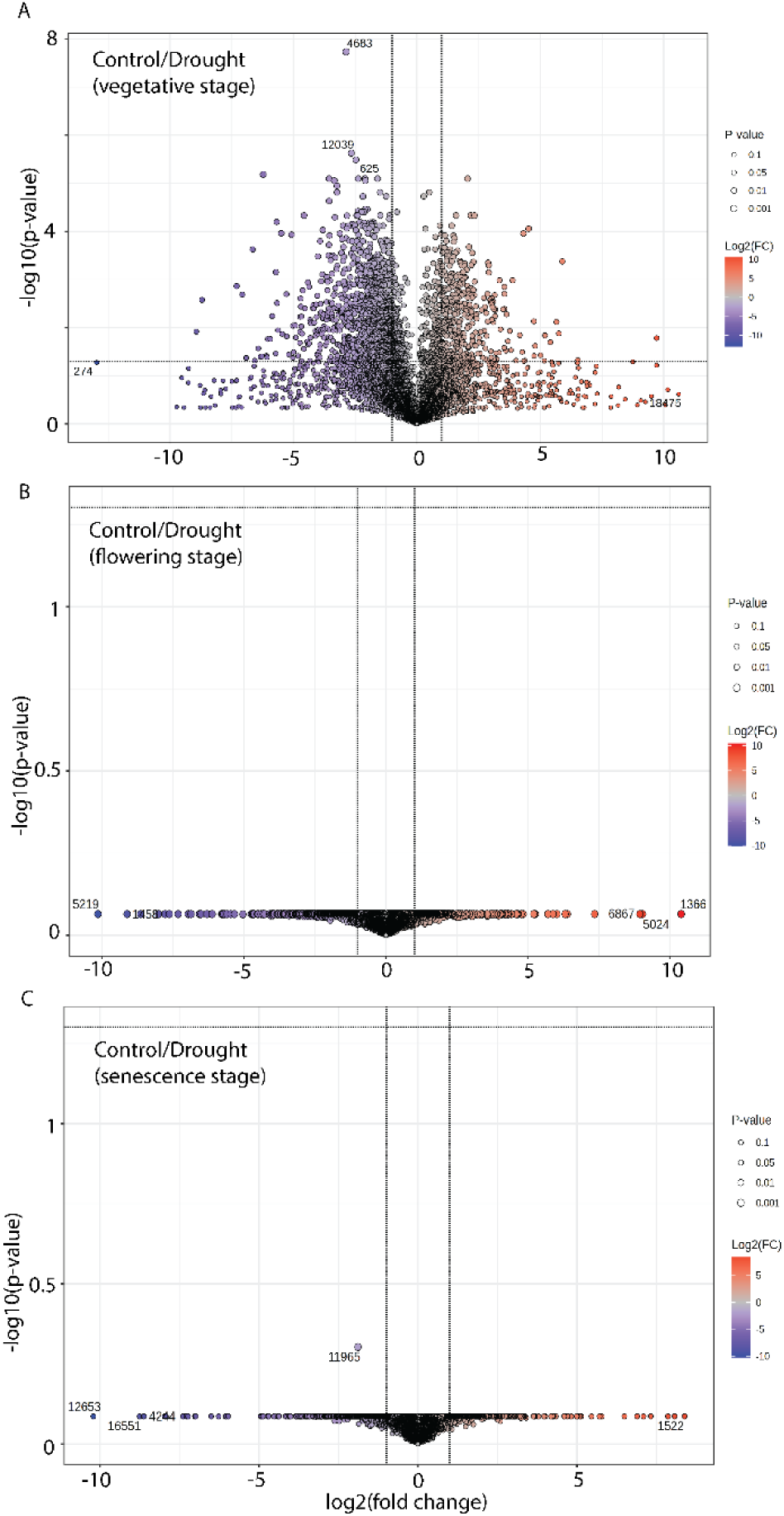
Volcano plots showing the differential accumulation of specialized metabolite features between Control and Drought treatments across three developmental stages. A. Vegetative stage, B. Flowering stage, and C. Senescence stage. Each point represents a metabolite, with significantly enriched metabolites in the Drought treatment shown in one orange and significantly depleted metabolites shown in purple. Metabolites that do not meet the significance threshold are represented in gray.

**Supplementary fig. 4.**
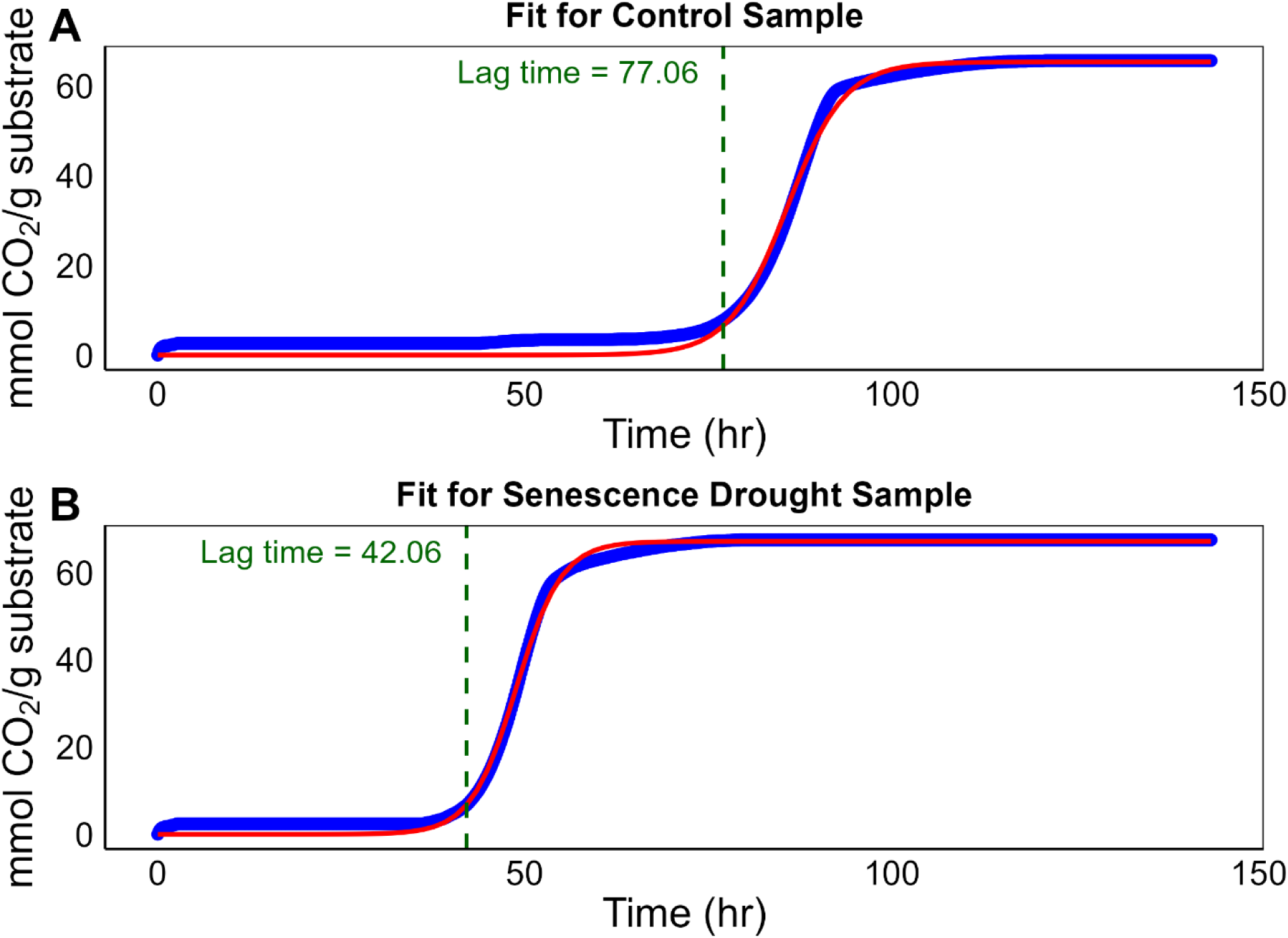
Exemplary CO₂ evolution curves showing substrate consumption rates over time. The x-axis represents time (hours), while the y-axis represents the substrate consumption rate (mmol CO₂/g substrate). The lag phase is defined as the time it takes for the fitted sigmoidal function to reach 10% of its maximum.

**Supplementary fig. 5.**
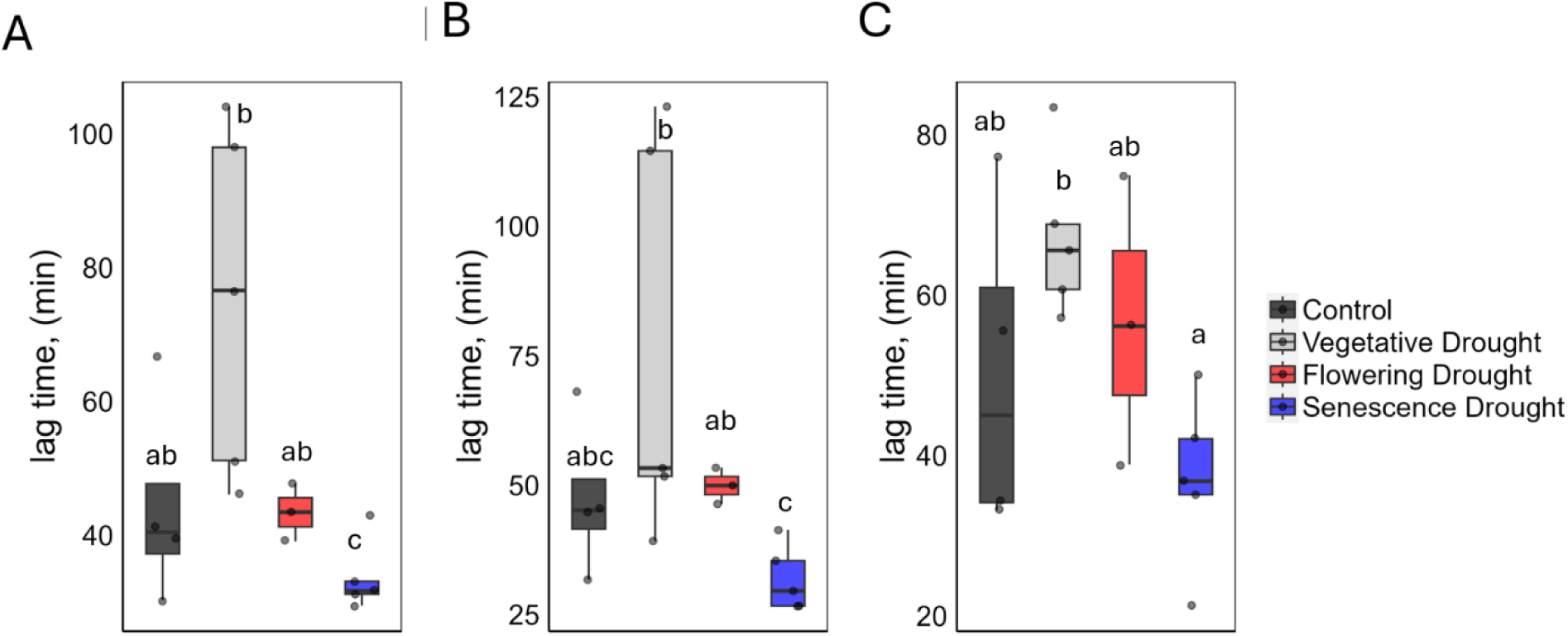
Lag phase of the CO_2_ evolution curve from three experiments (A) 1218 (B)1220 and (C) 1224. The lag phase was calculated by fitting CO_2_ evolution data to the sigmoidal curve:

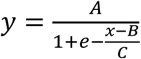 where A = maximum value of the curve (upper bound) x = time e = Euler’s number B = inflection point, where the curve changes shape C = rate/steepness of the curve.

**Supplementary fig.6.**
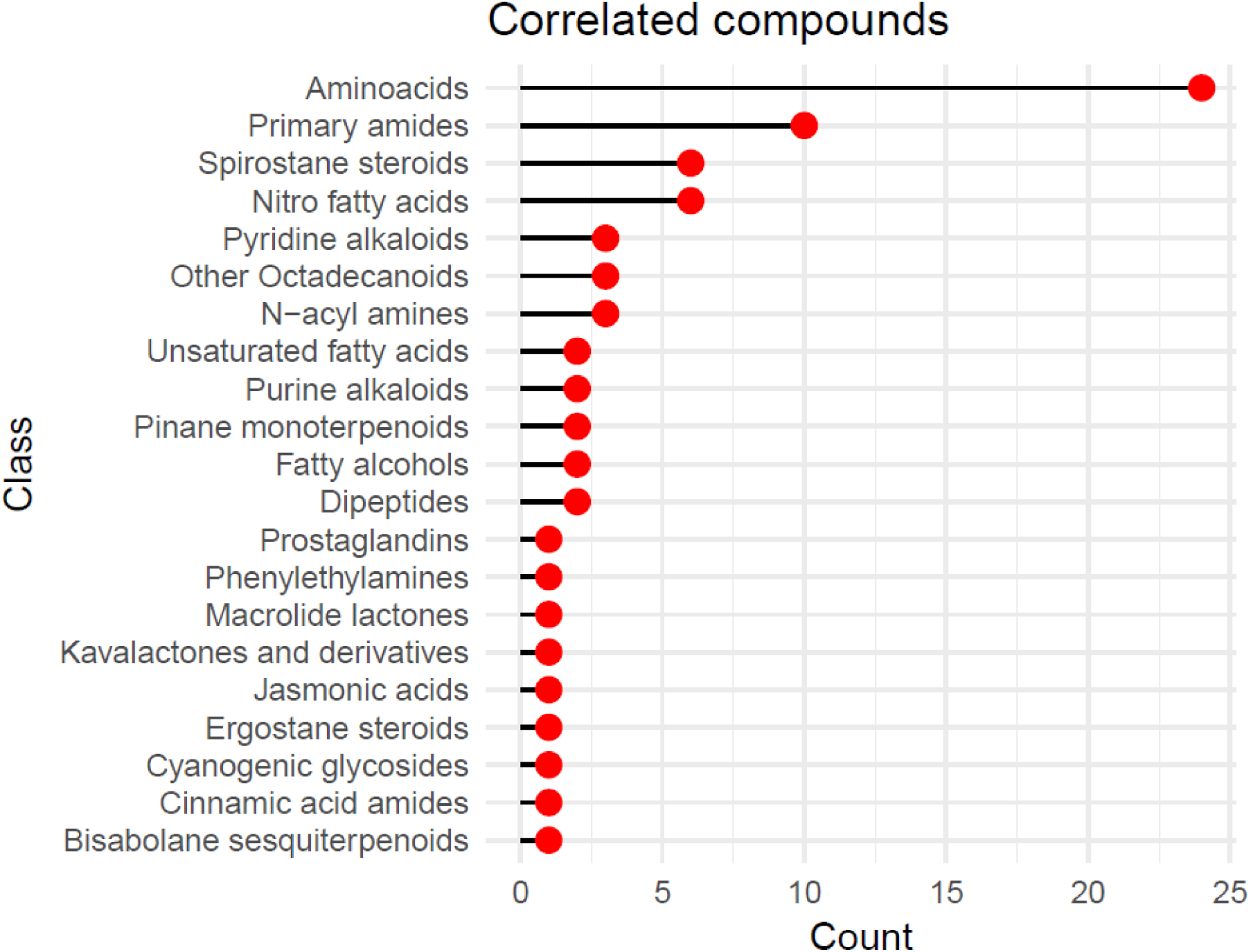
List of the chemical classes of metabolite features identified in the switchgrass hydrolysate by LC-MS that significantly correlated with fermentation lag time. X –axis shows the number of features within each compound groups

**Supplementary fig. 7.**
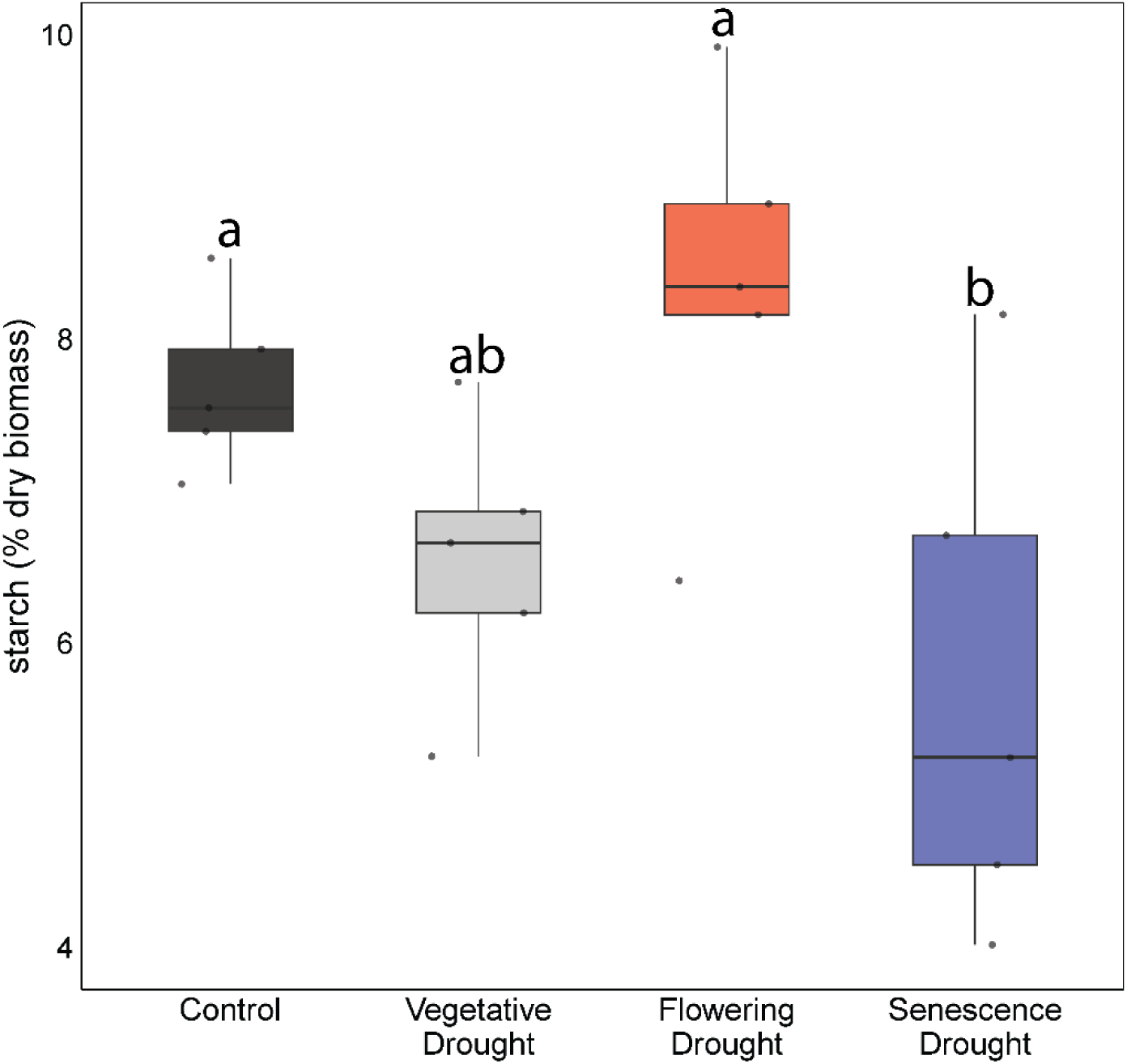
Total starch content expressed as a percentage of dry sample biomass. Bars show averages, N= 10, and vertical bars indicate standard error of mean. Groups with dissimilar letters are significantly different by Tukey’s post-hoc test (p<0.05).

**Supplementary fig. 8.**
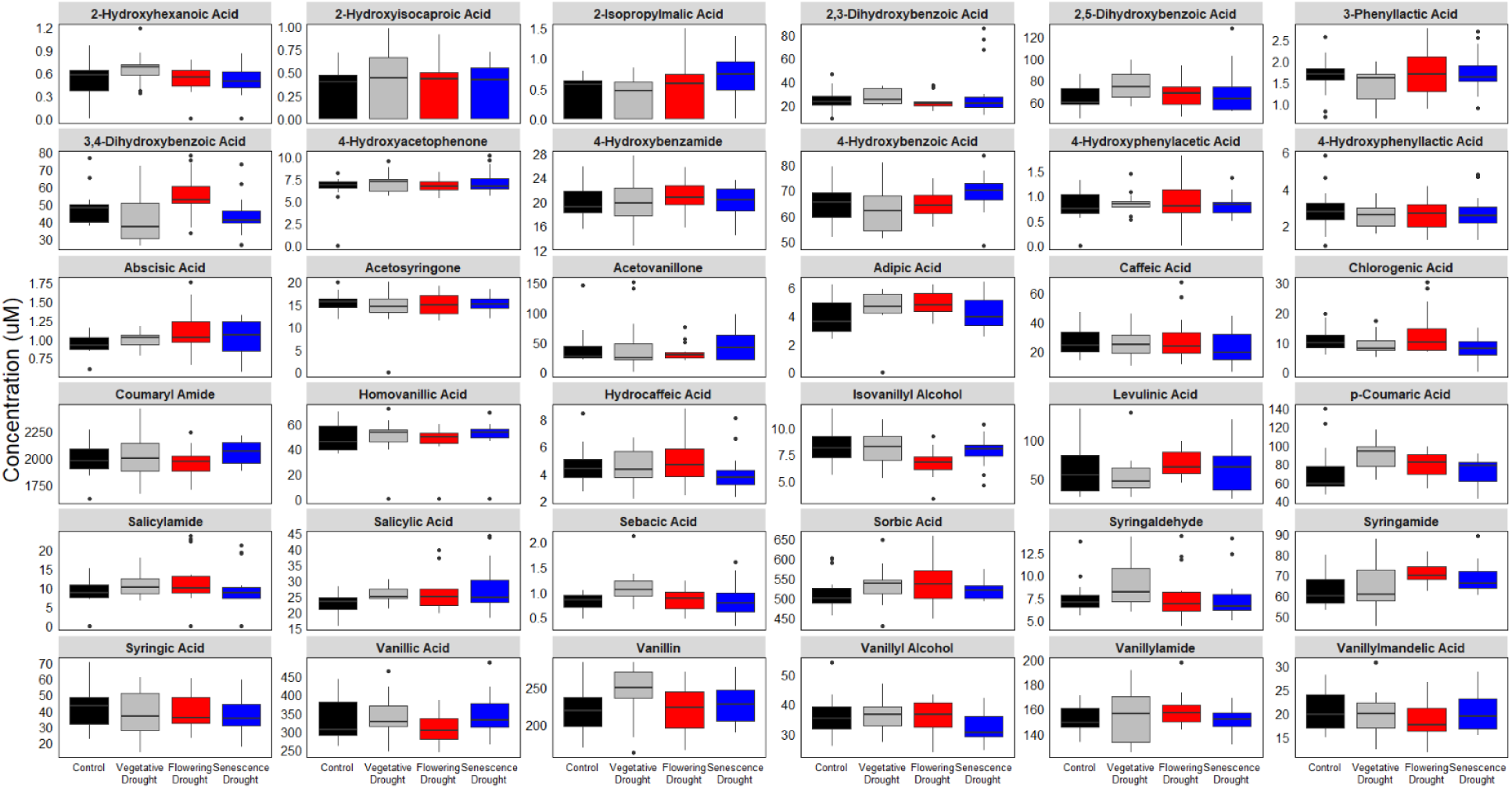
Concentration of switchgrass hydrolysate compounds that are not significantly different among treatment groups. The boxplot displays the distribution of concentration of hydrolysate compounds derived from lignocellulose and detected across different samples using LC-MS. Compounds were identified (N = 5) and quantified based on their retention time and m/z values, followed by data normalization and hierarchical clustering to reveal patterns of variation among samples. The boxes show the interquartile range (IQR, the middle 50% of values), the horizontal line inside the box represents the median, and the whiskers indicate the data spread. Outliers are displayed as individual points.

**Supplementary fig. 9.**
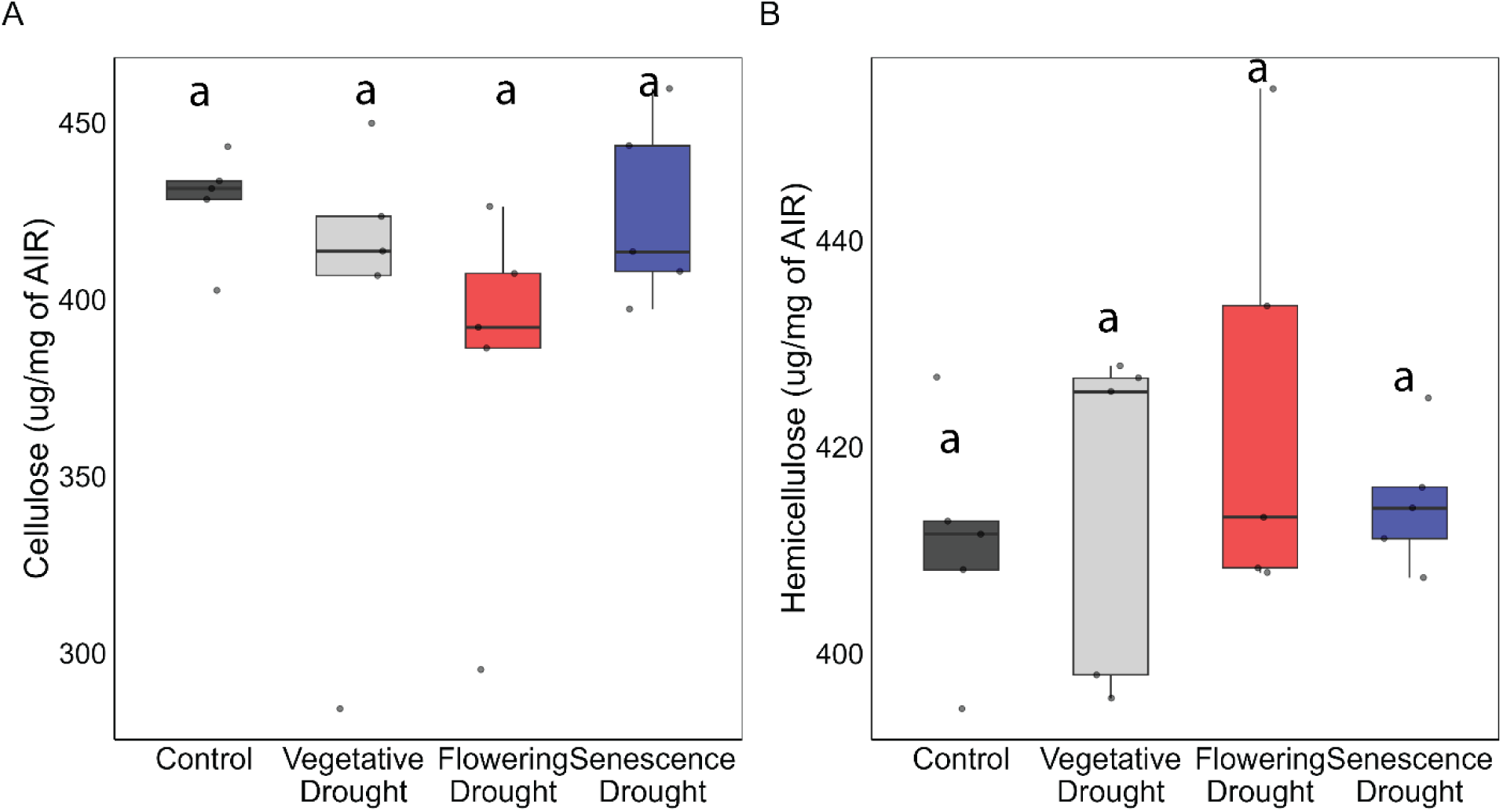
Matrix polysaccharides A. Cellulose and B. Hemicellulose content per milligram of alcohol insoluble residue (AIR) and B. The aboveground dry biomass samples were ground and homogenized to perform the fiber analysis. Bars show averages, N= 10, and vertical bars indicate standard error of mean. Groups with dissimilar letters are significantly different by Tukey’s post-hoc test (p<0.05).

**Supplementary fig. 10.**
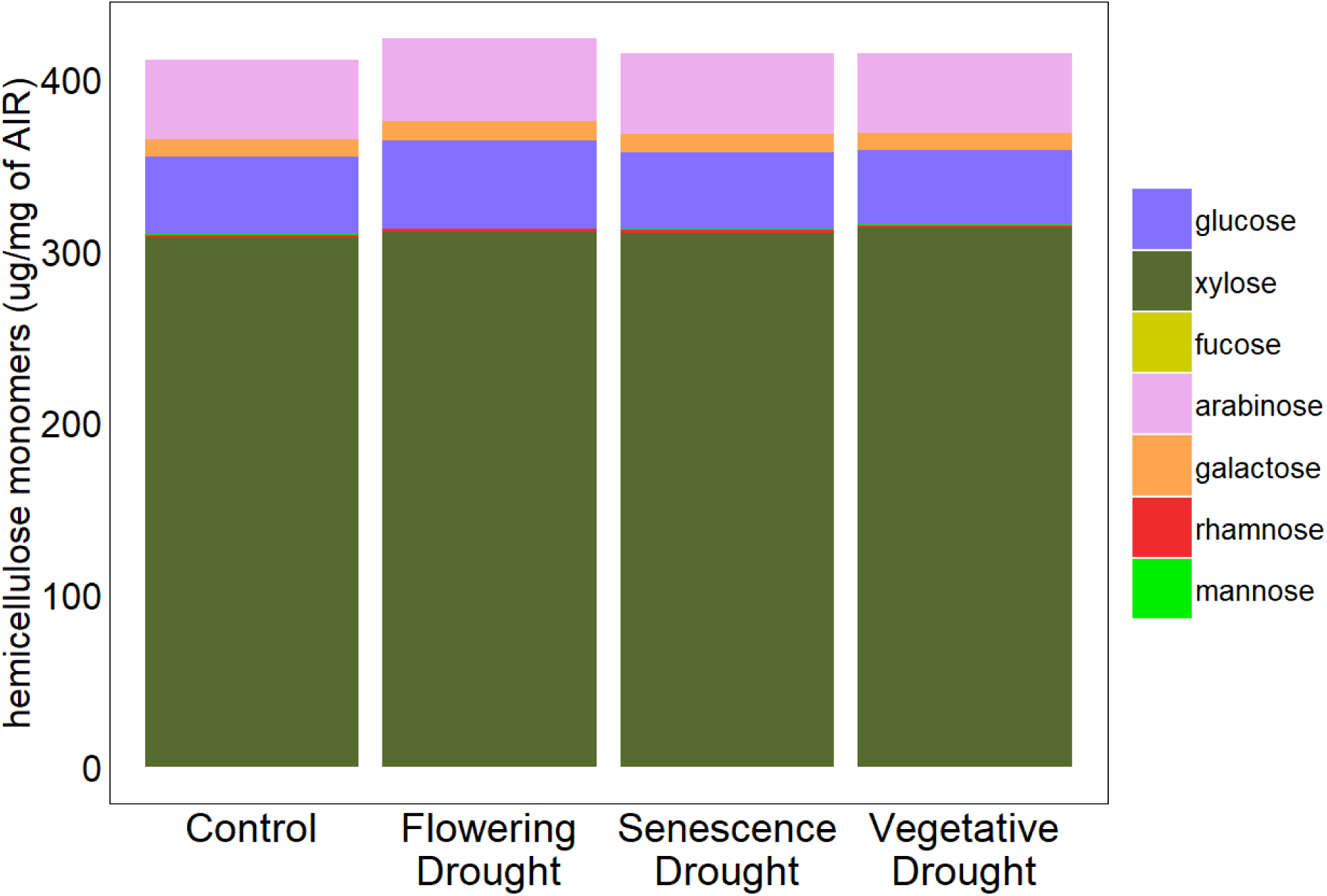
Monosaccharide components of hemicellulose per milligram of alcohol insoluble residue (AIR) and B. The aboveground dry biomass samples were ground and homogenized to perform fiber analysis. The sum total of the monomers gives total hemicellulose content. No significant difference between the treatments was detected in the monomers.

**Supplementary fig. 11.**
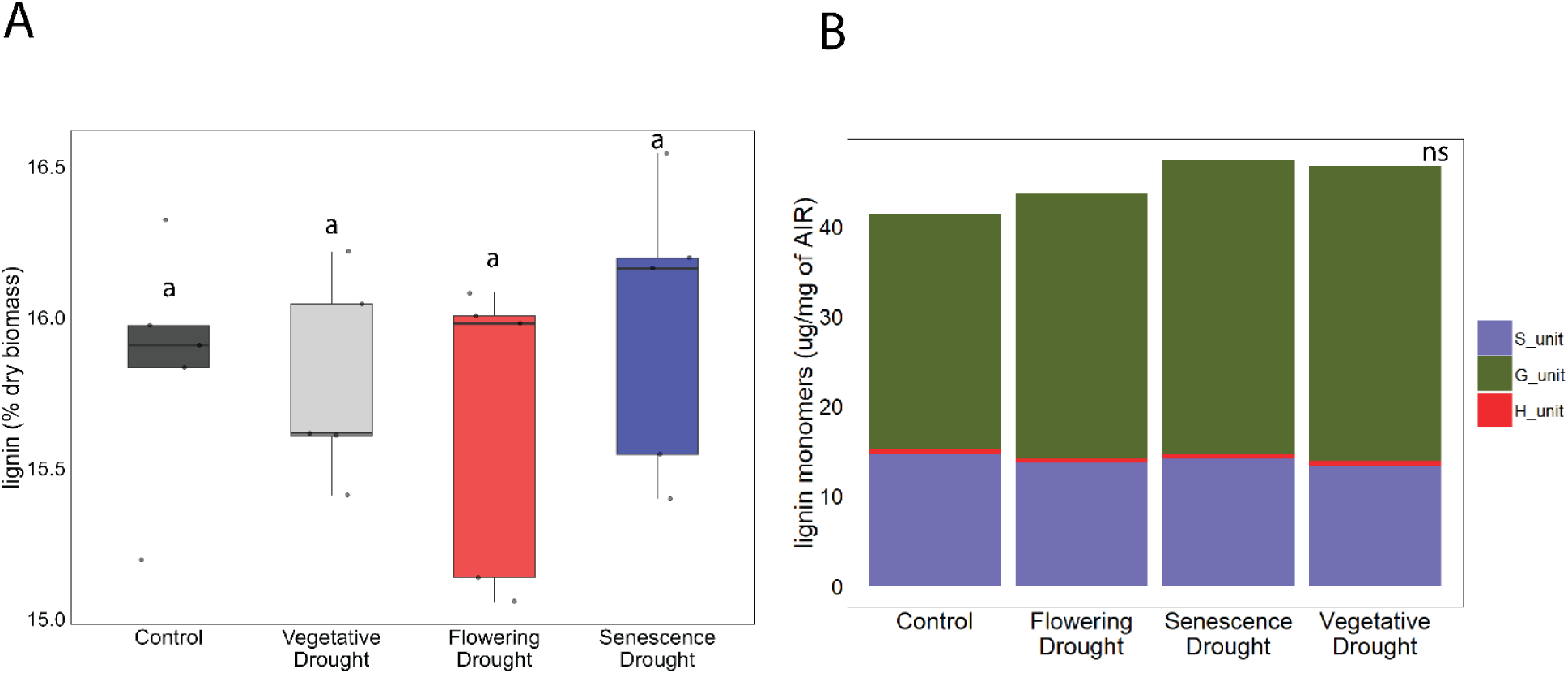
A. Total lignin and B. monolignol derived constituent lignin units (S Unit = Syringyl unit, G unit = Guaiacyl unit and H unit = *p*-hydroxyphenyl unit) expressed as % dry biomass and content per milligram of alcohol insoluble residue (AIR) respectively. The aboveground dry biomass samples were ground and homogenized to perform the fiber analysis. Bars show averages, N= 10, and vertical bars indicate standard error of mean. Groups with dissimilar letters are significantly different by Tukey’s post-hoc test (p<0.05). No significant difference between the treatments was detected in the lignin constituent units.

**Supplementary fig. 12.**
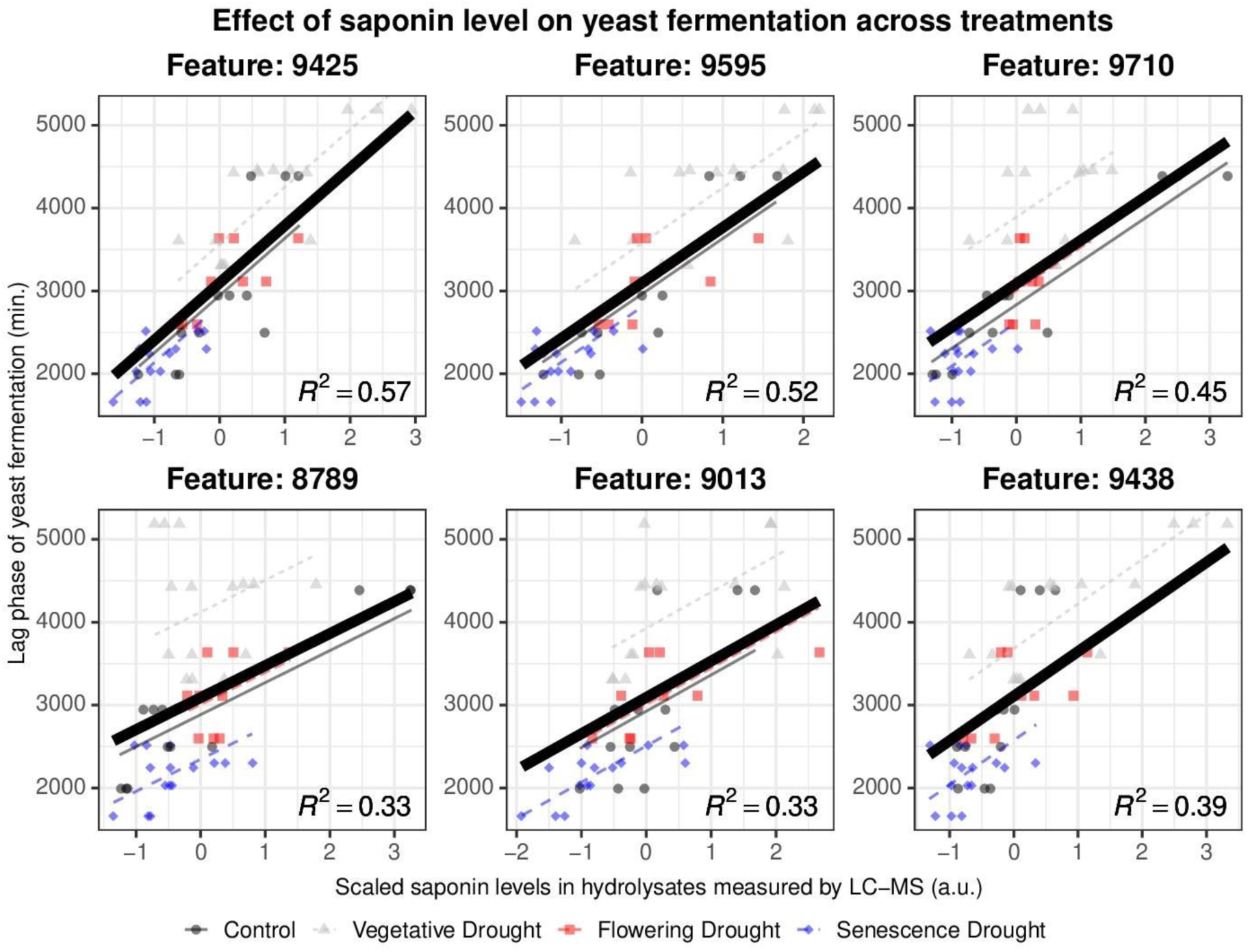
Scatter plot of lag time as a function of scaled saponin content, with linear regression lines shown per treatment group. Lines reflect fixed slopes from a mixed-effects model.

**Supplementary fig. 13.**
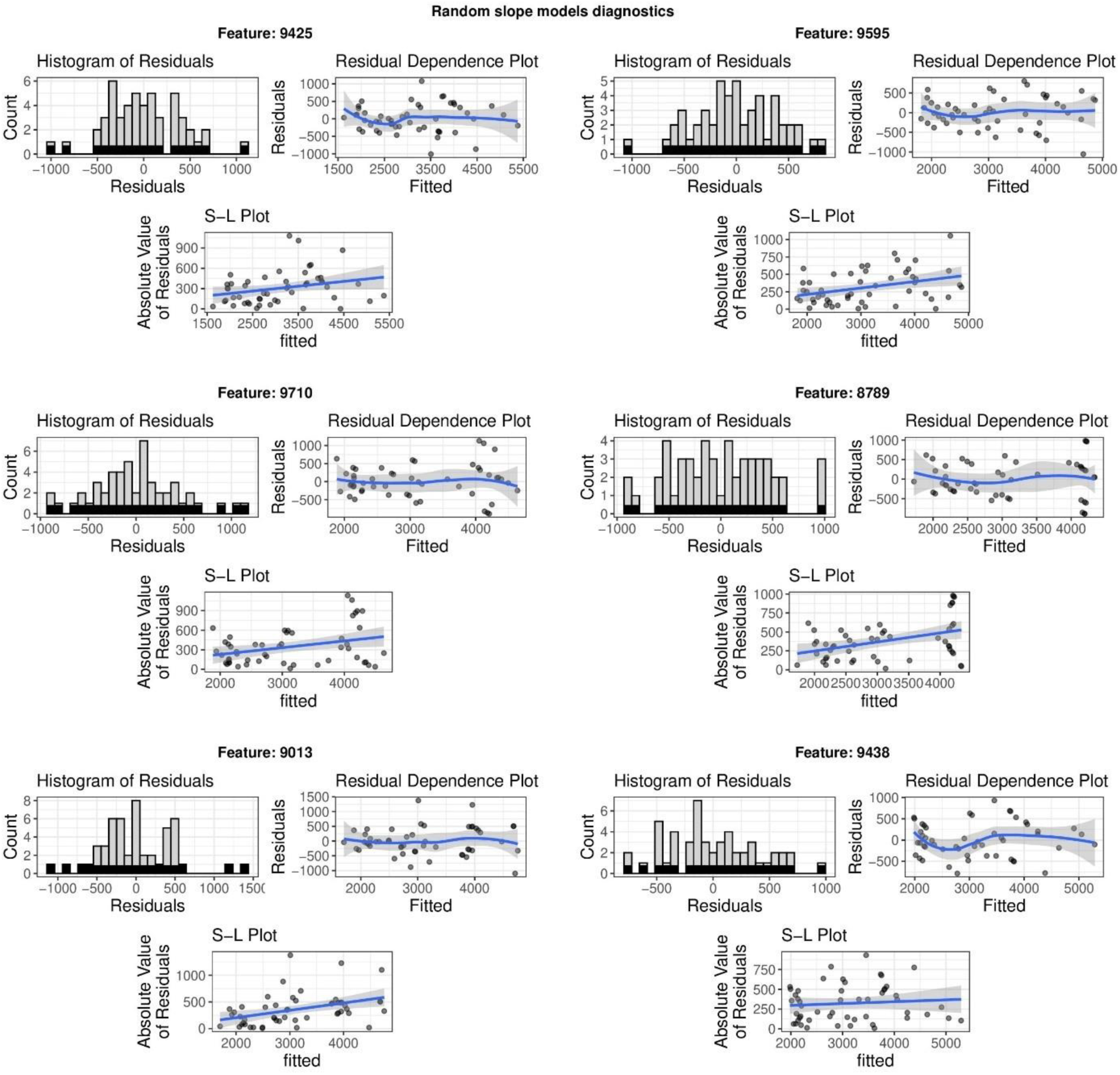
model diagnostics for models with random slope

**Supplementary fig. 14.**
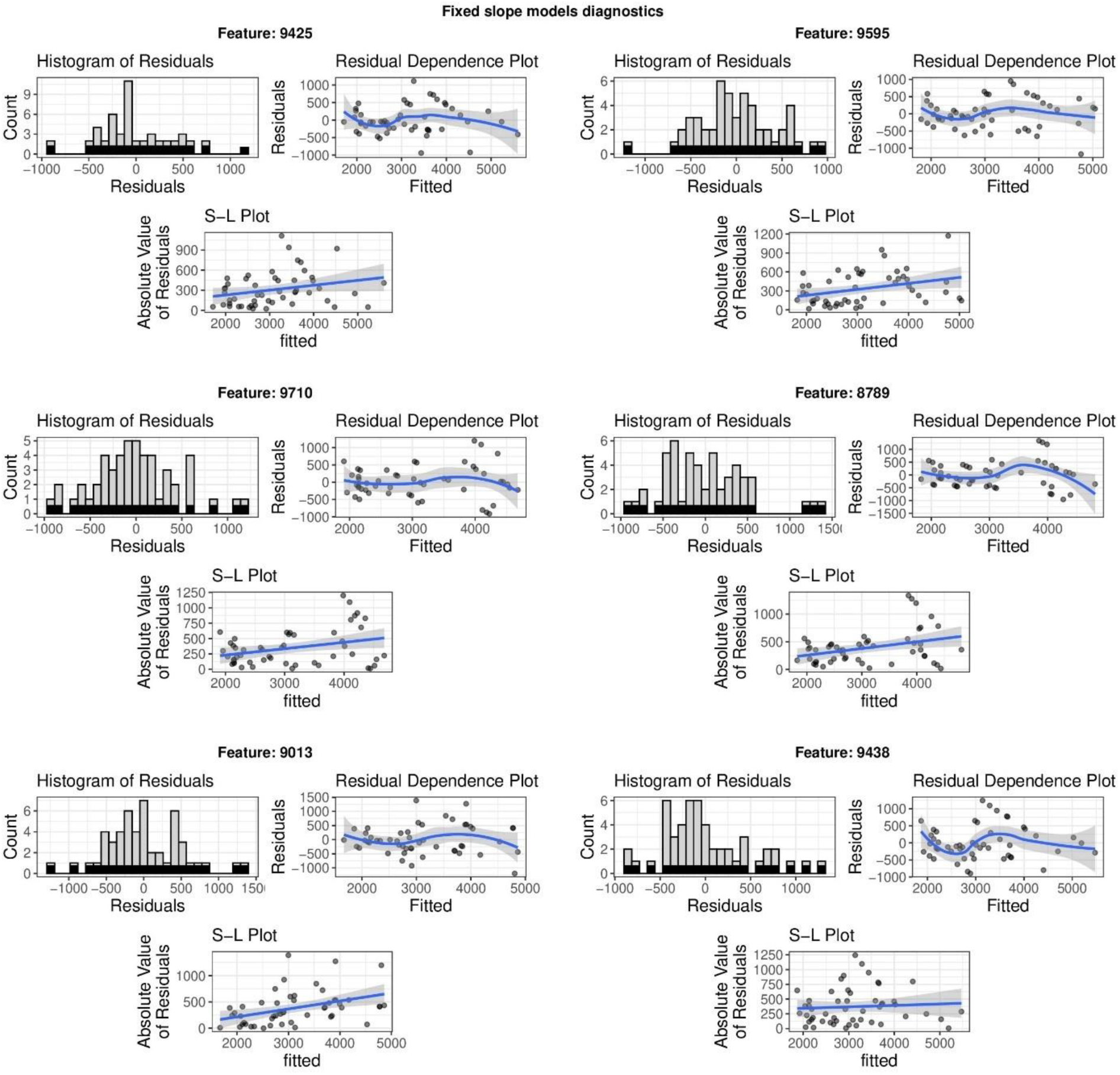
model diagnostics for model with fixed slope fermentation lag time

**Supplementary Table 1:** List of hydrolysate features that significantly correlated with fermentation lag time.

| Feature ID | correlation | cor_pvalue | Padj |
| --- | --- | --- | --- |
| 20 | 0.509364513 | 1.79472E-07 | 6.63404E-06 |
| 2801 | 0.568263055 | 5.78132E-09 | 5.2797E-07 |
| 2835 | 0.515657126 | 1.26453E-07 | 4.97009E-06 |
| 2986 | 0.555671724 | 1.2414E-08 | 9.40135E-07 |
| 2987 | 0.630453697 | 1.04545E-10 | 5.10582E-08 |
| 2988 | 0.583634579 | 2.22488E-09 | 2.93613E-07 |
| 2989 | 0.582221117 | 2.43154E-09 | 2.93613E-07 |
| 3066 | 0.534734324 | 4.2678E-08 | 2.07343E-06 |
| 3292 | 0.591268634 | 1.37226E-09 | 2.36714E-07 |
| 3293 | 0.585055651 | 2.0344E-09 | 2.87128E-07 |
| 3295 | 0.507992712 | 1.93604E-07 | 6.99E-06 |
| 3406 | 0.571166249 | 4.8362E-09 | 5.00547E-07 |
| 3408 | 0.547091776 | 2.07029E-08 | 1.2362E-06 |
| 3409 | 0.542183088 | 2.76464E-08 | 1.56077E-06 |
| 3719 | 0.515302236 | 1.28989E-07 | 5.00638E-06 |
| 4108 | 0.571776476 | 4.6576E-09 | 4.98684E-07 |
| 4134 | 0.586122747 | 1.9019E-09 | 2.87128E-07 |
| 4322 | 0.591641357 | 1.34005E-09 | 2.36714E-07 |
| 4443 | 0.535442338 | 4.09625E-08 | 2.05143E-06 |
| 4545 | 0.601636207 | 7.05114E-10 | 1.82448E-07 |
| 4640 | 0.534710275 | 4.27374E-08 | 2.07343E-06 |
| 4644 | 0.569582707 | 5.33133E-09 | 5.17305E-07 |
| 4645 | 0.554289361 | 1.34871E-08 | 9.97085E-07 |
| 4646 | 0.556521067 | 1.17963E-08 | 9.15689E-07 |
| 4665 | 0.506654912 | 2.08418E-07 | 7.43838E-06 |
| 4887 | 0.525911226 | 7.08561E-08 | 3.05567E-06 |
| 4912 | 0.568486581 | 5.70258E-09 | 5.2797E-07 |
| 4944 | 0.550753193 | 1.66588E-08 | 1.10054E-06 |
| 5424 | 0.54605735 | 2.20085E-08 | 1.26549E-06 |
| 5427 | 0.531088336 | 5.2675E-08 | 2.37038E-06 |
| 5619 | 0.538875256 | 3.3549E-08 | 1.79603E-06 |
| 5626 | 0.527995942 | 6.29022E-08 | 2.79016E-06 |
| 5782 | 0.565717563 | 6.756E-09 | 5.99354E-07 |
| 5833 | 0.632947016 | 8.82757E-11 | 5.10582E-08 |
| 5856 | 0.574787336 | 3.86622E-09 | 4.36728E-07 |
| 5951 | 0.527126219 | 6.61094E-08 | 2.89112E-06 |
| 5967 | 0.592758876 | 1.24785E-09 | 2.36714E-07 |
| 6339 | 0.532214839 | 4.93659E-08 | 2.25414E-06 |
| 6409 | 0.550091409 | 1.7328E-08 | 1.1209E-06 |
| 6468 | 0.5589466 | 1.01922E-08 | 8.19131E-07 |
| 6477 | 0.612334824 | 3.50614E-10 | 1.08866E-07 |
| 6499 | 0.533474161 | 4.59051E-08 | 2.15963E-06 |
| 6569 | 0.519024816 | 1.04671E-07 | 4.33339E-06 |
| 6661 | 0.605565665 | 5.46268E-10 | 1.54197E-07 |
| 6843 | 0.593595804 | 1.18288E-09 | 2.36714E-07 |
| 6992 | 0.55879076 | 1.02886E-08 | 8.19131E-07 |
| 7029 | 0.514890632 | 1.31992E-07 | 5.05968E-06 |
| 7109 | 0.504828285 | 2.30426E-07 | 7.9497E-06 |
| 7110 | 0.51138157 | 1.60488E-07 | 6.0038E-06 |
| 7393 | 0.536739434 | 3.79915E-08 | 1.99938E-06 |
| 7507 | 0.59697913 | 9.52244E-10 | 2.23975E-07 |
| 7509 | 0.626289373 | 1.38479E-10 | 5.10582E-08 |
| 7512 | 0.533822384 | 4.49904E-08 | 2.14916E-06 |
| 7513 | 0.55081019 | 1.66024E-08 | 1.10054E-06 |
| 7678 | 0.552116263 | 1.53587E-08 | 1.05975E-06 |
| 7824 | 0.505352257 | 2.23893E-07 | 7.89987E-06 |
| 7900 | 0.587455314 | 1.74821E-09 | 2.85694E-07 |
| 7908 | 0.553496286 | 1.41429E-08 | 1.017E-06 |
| 8139 | 0.532920873 | 4.73957E-08 | 2.19647E-06 |
| 8299 | 0.574489412 | 3.93829E-09 | 4.36728E-07 |
| 8348 | 0.53968711 | 3.19962E-08 | 1.77304E-06 |
| 8353 | 0.508887292 | 1.84271E-07 | 6.73131E-06 |
| 8354 | 0.547895377 | 1.97409E-08 | 1.22591E-06 |
| 8458 | 0.585508155 | 1.97715E-09 | 2.87128E-07 |
| 8461 | 0.536243364 | 3.91022E-08 | 2.02098E-06 |
| 8462 | 0.596064334 | 1.00987E-09 | 2.23975E-07 |
| 8563 | 0.50488705 | 2.29684E-07 | 7.9497E-06 |
| 8635 | 0.560984315 | 9.01033E-09 | 7.56137E-07 |
| 8845 | 0.627581325 | 1.26938E-10 | 5.10582E-08 |
| 9011 | 0.51847907 | 1.07936E-07 | 4.40975E-06 |
| 9265 | 0.515733692 | 1.25912E-07 | 4.97009E-06 |
| 9321 | 0.511771678 | 1.57048E-07 | 5.94675E-06 |
| 9423 | 0.504243005 | 2.37941E-07 | 8.11875E-06 |
| 9425 | 0.65553225 | 1.85241E-11 | 3.19987E-08 |
| 9426 | 0.646830748 | 3.40167E-11 | 3.19987E-08 |
| 9427 | 0.547402877 | 2.03252E-08 | 1.2362E-06 |
| 9438 | 0.57006411 | 5.17581E-09 | 5.17305E-07 |
| 9568 | 0.644056679 | 4.12221E-11 | 3.19987E-08 |
| 9569 | 0.522896408 | 8.41046E-08 | 3.57733E-06 |
| 9593 | 0.625300974 | 1.47995E-10 | 5.10582E-08 |
| 9595 | 0.647346795 | 3.28195E-11 | 3.19987E-08 |
| 9710 | 0.582506059 | 2.38842E-09 | 2.93613E-07 |
| 10140 | 0.546492553 | 2.14497E-08 | 1.25663E-06 |
| 10193 | 0.535980405 | 3.97037E-08 | 2.02098E-06 |
| 10194 | 0.539394049 | 3.25485E-08 | 1.77304E-06 |
| 10768 | 0.582044757 | 2.4586E-09 | 2.93613E-07 |
| 10953 | 0.553181612 | 1.44116E-08 | 1.017E-06 |
| 11063 | 0.517685223 | 1.12862E-07 | 4.55111E-06 |
| 11100 | 0.522465927 | 8.6182E-08 | 3.61615E-06 |
| 11120 | 0.561263844 | 8.85897E-09 | 7.56137E-07 |
| 11182 | 0.548329835 | 1.9239E-08 | 1.21912E-06 |

**Supplementary Table 2.** Macro and micro-nutrient profile of the potting media.

|  |  |
| --- | --- |
| Total Nitrogen | 0.27% |
| Available Phosphate | 0.23% |
| Soluble Potash (K2O) | 0.90% |
| Calcium | 0.22% |
| Magnesium | 0.11% |
| Sulfur | 0.30% |
| Boron | 0.008% |
| Copper | 0.02% |
| Iron | 0.42% |
| Manganese | 0.07% |
| Molybdenum | 0.007% |
| Zinc | 0.07% |

**Supplementary table 3.** Model estimates with corresponding p values.

| Random intercept and fixed slope models |  |  |  |  |  |  |  |
| --- | --- | --- | --- | --- | --- | --- | --- |
| Saponin | Estimate | Std.Error | df | t | value | Pr(>\|t\|) | Signif |
| 9425 | (Intercept) | 3098.22 | 183.75 | 2.73 | 16.86 | 0.000770 | *** |
| 9425 | Mass_scaled | 691.6 | 79.37 | 48.66 | 8.71 | 0.000000 | *** |
| 9595 | (Intercept) | 3097.8 | 190.2 | 2.7 | 16.29 | 0.000900 | *** |
| 9595 | saponin_scaled | 668.7 | 83.9 | 48.5 | 7.97 | 0.000000 | *** |
| 9710 | (Intercept) | 3087.74 | 296.42 | 2.99 | 10.42 | 0.001900 | ** |
| 9710 | saponin_scaled | 523.76 | 80.47 | 47.66 | 6.51 | 0.000000 | *** |
| 8789 | (Intercept) | 3090.57 | 387.1 | 3.03 | 7.98 | 0.003900 | ** |
| 8789 | saponin_scaled | 386.6 | 76.91 | 46.33 | 5.03 | 0.000008 | *** |
| 9013 | (Intercept) | 3092.83 | 317.42 | 2.98 | 9.74 | 0.002300 | ** |
| 9013 | saponin_scaled | 438.04 | 85.67 | 47.4 | 5.11 | 0.000006 | *** |
| 9438 | (Intercept) | 3108.91 | 248.43 | 2.83 | 12.51 | 0.001500 | ** |
| 9438 | saponin_scaled | 537.81 | 89.15 | 48.78 | 6.03 | 0.000000 | *** |
| Random intercept and random slope models |  |  |  |  |  |  |  |
| Saponin | Estimate | Std.Error | df | t | value | Pr(>\|t\|) | Signif |
| 9425 | (Intercept) | 3149.39 | 162.95 | 3.68 | 19.33 | 0.0000780 | *** |
| 9425 | saponin_scaled | 764.85 | 86.45 | 7.37 | 8.85 | 0.0000350 | *** |
| 9595 | (Intercept) | 3156.22 | 173.78 | 3.59 | 18.16 | 0.0001200 | *** |
| 9595 | saponin_scaled | 723.74 | 93.22 | 7.16 | 7.76 | 0.0000980 | *** |
| 9710 | (Intercept) | 3120.46 | 269.84 | 3.88 | 11.56 | 0.0003800 | *** |
| 9710 | saponin_scaled | 503.75 | 90.27 | 5.18 | 5.58 | 0.0022800 | ** |
| 8789 | (Intercept) | 3118.48 | 334.71 | 4.06 | 9.32 | 0.0006800 | *** |
| 8789 | saponin_scaled | 346.07 | 140.8 | 3.28 | 2.46 | 0.0837900 | . |
| 9013 | (Intercept) | 3102.99 | 282.34 | 3.94 | 10.99 | 0.0004200 | *** |
| 9013 | saponin_scaled | 462.46 | 111.4 | 2.64 | 4.15 | 0.0324000 | * |
| 9438 | (Intercept) | 3144.04 | 257.18 | 3.63 | 12.22 | 0.0004500 | *** |
| 9438 | saponin_scaled | 664.87 | 252.81 | 3.25 | 2.63 | 0.0720000 | . |

## Notes

### Competing Interest Statement

The authors have declared no competing interest.

